# Human RPL7 and DDX21 interact with HTLV-1 Gag and enhance tRNA^Pro^ primer annealing to genomic RNA

**DOI:** 10.1101/2025.07.15.664966

**Authors:** Yu-Ci Syu, Zixi Long, Karin Musier-Forsyth

## Abstract

Human T-cell leukemia virus type 1 (HTLV-1), an oncogenic retrovirus, uses human tRNA^Pro^ to prime reverse transcription (RT). However, how tRNA^Pro^ is annealed to the primer binding site (PBS), which is embedded in a highly structured hairpin in the genomic RNA (gRNA), remains unclear. We hypothesize that HTLV-1 Gag may have more robust chaperone activity than mature HTLV-1 nucleocapsid (NC), which in contrast to HIV-1 NC, displays relatively weak chaperone function, and that a cellular co-factor may be required to facilitate primer tRNA annealing. Recombinant HTLV-1 Gag was successfully purified for the first time and used to perform primer-annealing assays. Relative to mature NC and matrix (MA) domains, HTLV-1 Gag is only slightly more effective at chaperoning the annealing of tRNA^Pro^ to the PBS. To identify potential HTLV-1 Gag interacting co-chaperones of tRNA annealing in cells, we performed affinity tagging/purification-mass spectrometry (AP-MS). Two significant AP-MS hits, RPL7 and DDX21, were further validated by reciprocal co-IP studies in both HEK293T and chronically HTLV-1-infected MT-2 cells. Domain mapping studies revealed that HTLV-1 Gag interacts with RPL7 and DDX21 through the zinc fingers in the NC domain independent of the presence of RNA. In addition, we showed that both RPL7 and DDX21 are packaged into virions. RPL7 or DDX21 alone was more effective than HTLV-1 Gag at annealing tRNA^Pro^ to the PBS. Synergistic effects of the Gag/RPL7/DDX21 combination in facilitating tRNA^Pro^ annealing to the PBS were found. Taken together, the mechanistic insights gained from these studies could be exploited for the development of new therapeutic strategies aimed at targeting HTLV-1 RT.

**Highlights:**

1. Mechanism for HTLV-1 reverse transcription primer annealing to the primer binding site in viral RNA is proposed.
2. Recombinant HTLV-1 Gag was purified successfully from *E. coli* for the first time.
3. HTLV-1 Gag interacts directly with human RPL7 and DDX21 in cells.
4. RPL7 and DDX21 enhance HTLV-1 Gag’s tRNA^Pro^ primer annealing activity.
5. tRNA^Pro^ annealing activity: DDX21 + RPL7 + Gag > DDX21 + RPL7 > DDX21 > RPL7 > Gag.

## Introduction

All retroviruses use host cell tRNAs as primers to initiate reverse transcription (RT) (1). The 3’ 18-nucleotides (nt) of the primer tRNA are perfectly complementary to a region in the 5’ untranslated region (UTR) of the genomic RNA (gRNA) known as the primer-binding site (PBS). The mechanism of how the human immunodeficiency virus type 1 (HIV-1) RT primer, tRNA^Lys3^, is selectively packaged into virions and annealed by the NC domain of Gag to the HIV-1 PBS has been extensively studied (2–5). HIV-1 NC is a robust nucleic acid (NA) chaperone protein (6,7), displaying duplex destabilization, nucleic acid aggregation, as well as rapid on-off binding kinetics (7–10). Almost every step in RT, including primer annealing, extension, and multiple strand-transfer steps, requires either Gag or NC’s RNA chaperone activities, which facilitate NA rearrangements and conformational changes (3,11–13). The NC domain of HIV-1 Gag is also the major player in gRNA packaging; NC has a preference for binding to single-stranded G residues that are exposed in the packaging signal (Psi) located in the HIV-1 5’ UTR (14). In contrast, MA demonstrates stronger RNA chaperone activity and plays a more prominent role in specific Psi RNA binding than NC in deltaretroviruses, including bovine leukemia virus (BLV), HTLV-1 and HTLV-2, (15–17). In our previous study, we found that HTLV-1 uses a specific tRNA^Pro^ isodecoder as the RT primer (18). However, the viral and/or host factors required for tRNA^Pro^ annealing to the highly structured HTLV-1 PBS are still unknown.

The HTLV-1 PBS is complementary to 3’ 18-nt of human tRNA^Pro^. RNA-structure probing data showed that the HTLV-1 PBS is embedded in a highly stable hairpin containing 10 Watson-Crick base pairs (17), in contrast to the HIV-1 PBS, which is much less structured (19). The NC domain of HIV-1 Gag is a robust NA chaperone protein capable of facilitating human tRNA^Lys3^ annealing to the HIV-1 PBS in the absence of any other co-factors (3,7). HTLV-1 NC is a relatively weak chaperone protein due to the unique acidic C-terminal extension (7,20), and we hypothesize that HTLV-1 MA or full-length Gag may be required to facilitate primer tRNA^Pro^ annealing. Alternatively, a host cell co-factor may be required.

In HIV-1-infected cells, ribosomal protein L7 (RPL7) was shown to interact with the NC domain of HIV-1 Gag and be packaged into HIV-1 virions. RPL7 had a positive synergistic effect on the chaperone activity of HIV-1 Gag; adding both proteins together increased annealing of tRNA^Lys3^ to the HIV-1 PBS (21,22). Another host factor, RNA helicase A (RHA/DHX9), has also been shown to be involved in HIV-1 RT. RHA interacts with HIV-1 Gag and is packaged into virions (23). RHA, together with Gag, has been proposed to change the structure of gRNA to facilitate tRNA^Lys3^ annealing to the HIV-1 PBS (24). In addition to facilitating the primer annealing step, RHA has been shown to increase the elongation processivity of HIV-1 reverse transcriptase (25).

In this work, we investigated both viral and host cell factors as potential chaperone proteins capable of facilitating primer annealing to the highly structured HTLV-1 PBS. We successfully expressed and purified recombinant HTLV-1 Gag from *Escherichia coli* for the first time and characterized the HTLV-1 Gag interactome in human cells. We identified HTLV-1 Gag-interacting partners, RPL7 and DDX21, which are known chaperone proteins or helicases, respectively, and investigated their roles in regulating RT. Their interactions with HTLV-1 Gag were mapped in cells and their ability to facilitate tRNA^Pro^ annealing to the PBS was examined *in vitro*. These data suggest that HTLV-1 Gag recruits one or more cellular co-factors to ensure primer placement onto the PBS prior to initiation of RT. Disrupting these direct Gag/co-factor interactions may serve as a new anti-HTLV-1 therapeutic strategy.

## Results

### Recombinant HTLV-1 Gag was successfully purified from *E. coli* for the first time and shown to be monomeric in solution

To investigate potential proteins responsible for annealing the HTLV-1 RT primer, tRNA^Pro^, to the highly structured HTLV-1 PBS (17), annealing assays were initially performed using viral chaperone protein candidates, such as HTLV-1 MA, HTLV-1 NC, and HIV-1 NC. The *in vitro* transcribed human tRNA^Pro^ and viral genome-derived PBS domain used in this study are shown in Figure 1A. Neither HTLV-1 MA, NC, nor the robust chaperone protein, HIV-1 NC (7), could facilitate the annealing of tRNA^Pro^ to the stable PBS domain *in vitro* (data not shown).

**Figure 1.**
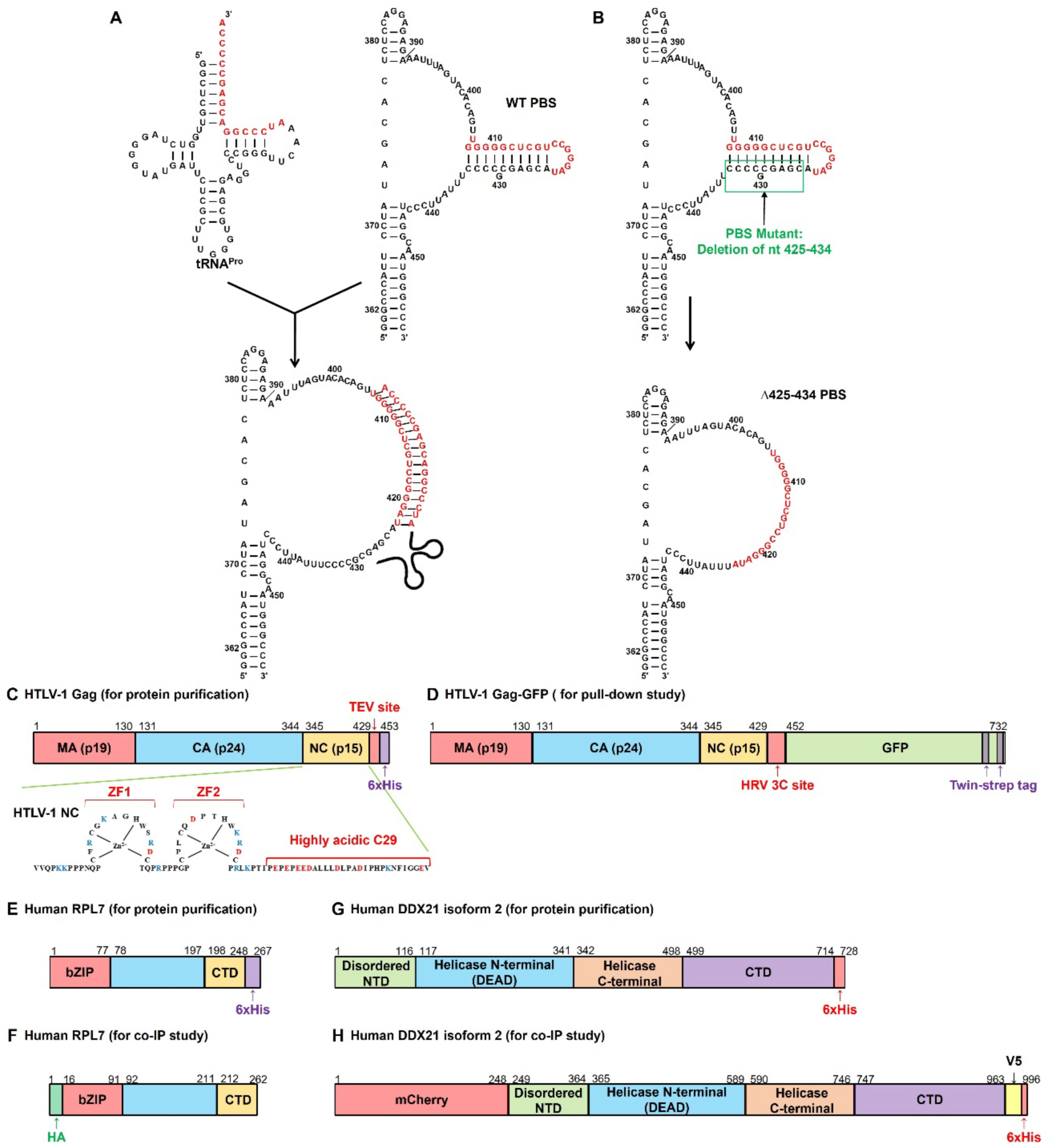
Schematic representation of RNAs and proteins used in this study. (A) Sequences and secondary structures of tRNA^Pro^ (75 nt, top left) and a portion of HTLV-1 5’ UTR containing the primer binding site (PBS) (98 nt, WT PBS, top right) used in this work. The sequence in the HTLV-1 PBS complementary to the 3’ 18-nt of tRNA^Pro^ is indicated by red letters. The bottom of this panel shows the tRNA^Pro^-PBS annealed complex. The structure of the PBS region is based on the structure-probing results reported in Wu et al (17). (B) Sequence and secondary structure of HTLV-1 PBS region before (WT PBS, top panel) and after deleting nt 425-434 (88 nt, Δ452-434 PBS mutant, bottom of this panel). (C) Domain architecture of HTLV-1 Gag bacterial expression construct consisting of matrix (MA, p19), capsid (CA, p24), and nucleocapsid (NC, p17) domains. The Tobacco Etch Virus (TEV) protease cleavage site and a 6xHis-tag at the C-terminus are indicated. The sequence of amino acids in the NC domain is shown. (D) Domain organization of HTLV-1 Gag mammalian expression construct used in AP-MS pull-down study; the human rhinovirus (HRV) 3C protease cleavage site, green fluorescent protein (GFP), and twin-strep tag at the C-terminus are indicated. (E) Domain organization of human RPL7 bacterial expression construct with a 6xHis-tag at the C-terminus, an N-terminal basic leucine zipper (bZIP), and a C-terminal domain (CTD). (F) Domain organization of human RPL7 mammalian expression construct with HA tag at the N-terminus. (G) Domain organization of human DDX21 bacterial expression construct with a 6xHis-tag at the C-terminus, a disordered N-terminal domain (NTD), a helicase core (the helicase N-terminal and C-terminal domains), and a CTD. (H) Human DDX21 mammalian expression construct with mCherry at the N-terminus and V5 and 6xHis tags at the C-terminus.

Because the mature NC or MA domains of HTLV-1 Gag failed to facilitate tRNA^Pro^ annealing to the PBS, we hypothesized that full-length HTLV-1 Gag may have more robust chaperone activity and facilitate tRNA^Pro^ annealing. To test this hypothesis, we successfully purified HTLV-1 Gag from *E. coli* and characterized its oligomeric state and chaperone activity *in vitro*. In our previous study, removing the C-terminal 29 highly acidic residues in HTLV-1 NC increased HTLV-1 NC’s chaperone activity, making it comparable to other retroviral NC proteins (20). To test whether deleting the C-terminal 29 residues in the NC domain of HTLV-1 Gag impacts its chaperone activity, HTLV-1 ΔC29 Gag was also purified (Figure 1C). The purity of the recombinant proteins was high, as shown by SDS-PAGE and Coomassie blue staining (Figure S1A).

Gag proteins play important roles in packaging gRNA and assembling viral particles. In HIV-1-infected cells, lower-order Gag oligomers, such as dimers or trimers, are formed in the cytoplasm, and higher-order Gag multimers develop at the plasma membrane (PM) (26–28). In contrast to HIV-1, there are no prominent HTLV-1 Gag-Gag interactions in the cytoplasm, and Gag oligomerization happens exclusively at the PM (27–30). To investigate the oligomeric states of purified HTLV-1 WT and ΔC29 Gag proteins in solution, size-exclusion chromatography with multi-angle laser-light scattering (SEC-MALS) analysis was performed. In the spectra shown in Figure S1B, only one peak was observed for each protein corresponding to calculated molecular weights of 59.6 kD for WT and 56 kD for ΔC29. Based on the theoretical molecular weights of 48.4 kD and 45.3 kD for WT and ΔC29, respectively, these results indicate that HTLV-1 Gag proteins form monomers in solution, consistent with the previous findings (27,28).

### HTLV-1 WT and ΔC29 Gag show weak primer annealing activity

To evaluate the annealing activities of HTLV-1 Gag proteins, tRNA^Pro^-PBS annealing assays were performed. In the annealing assays, 5’ ^32^P-labeled tRNA^Pro^ and 10-fold excess of the HTLV-1 5’ UTR PBS domain (98 nt) were folded separately and then incubated together in the presence of different concentrations of proteins at 37°C (Figure 1A). In concentration-dependence annealing assays (Figure 2A - 2D), tRNA^Pro^ was annealed to the HTLV-1 PBS in the presence of varying concentrations of HTLV-1 WT or ΔC29 Gag at 37°C for 1 h. The results are shown in Figure 2A, the lowest band is the free tRNA^Pro^ (F form) and the major shifted bands (B1 and B2) likely represent two different conformations of the tRNA^Pro^-PBS complex. Even in the heat-annealing positive control, very little annealed complex is formed. Although very little complex formation was observed in the presence of HTLV-1 WT or ΔC29 Gag, deletion of the C-terminal 29 acidic residues improved HTLV-1 Gag’s chaperone activity, as predicted. In the presence of 4 μM HTLV-1 WT Gag, only 5.91 ± 1.2% of tRNA^Pro^ is annealed to the PBS. K_1/2_ for WT Gag is 1.99 ± 1.6 μM, and the catalytic efficiency (Annealed_max_/K_1/2_) is 2.97 ± 2.5 μM^-1^ (Figure 2A and 2C; Table 1). In contrast, with 4 μM HTLV-1 ΔC29 Gag, 25.7 ± 7.3% of tRNA^Pro^ is annealed to the PBS. K_1/2_ for Gag ΔC29 is 4.03 ± 2.6 μM, and the catalytic efficiency is 6.38 ± 4.5 μM^-1^ (Figure 2A and 2C; Table 1). Based on these data, HTLV-1 ΔC29 Gag chaperoned tRNA^Pro^ annealing to the PBS 2.1-fold more efficiently than WT (Table 1).

**Figure 2.**
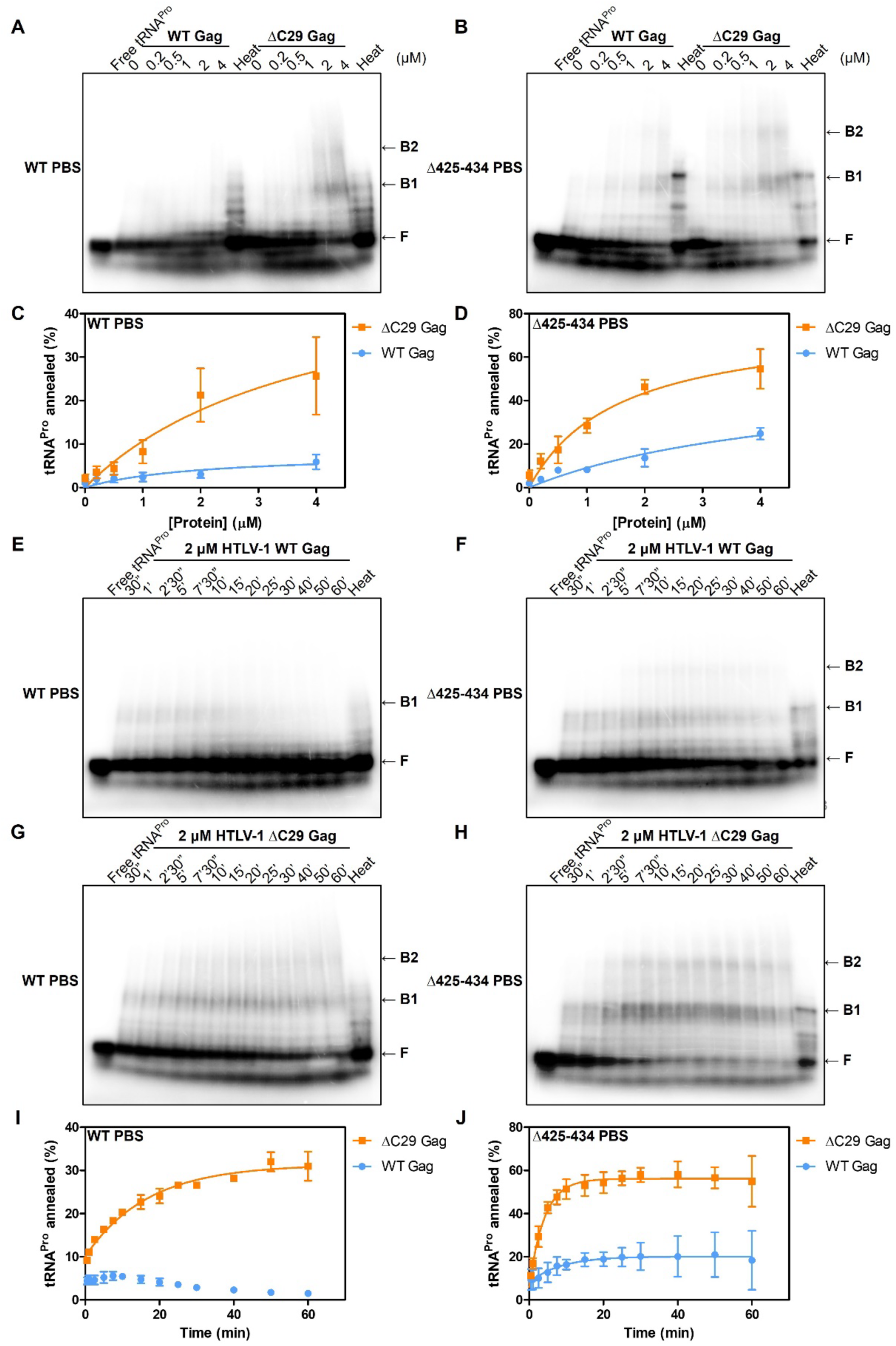
Concentration-dependence and time-course annealing assays show that HTLV-1 ΔC29 Gag chaperones tRNA^Pro^ annealing to the stable HTLV-1 WT PBS and the less structured HTLV-1 Δ425-434 PBS more effectively than HTLV-1 WT Gag. (A and B): Concentration-dependence annealing assays using 20 nM 5’ ^32^P-labeled tRNA^Pro^ and 200 nM HTLV-1 WT PBS (A) or Δ425-434 PBS (B) in the presence of varying concentrations of HTLV-1 WT or ΔC29 Gag at 37°C for 1 h. F indicates free tRNA^Pro^ and B1 and B2 indicate different conformations of tRNA^Pro^-PBS binary complexes. Heat indicates heat annealing, the positive control. (C and D): Graphs showing percentages of tRNA^Pro^ annealed to WT (C) or Δ425-434 PBS (D) in the presence of varying concentrations of HTLV-1 WT or ΔC29 Gag. (E-H): Time-course annealing assays using 20 nM 5’ ^32^P-labeled tRNA^Pro^ and 200 nM HTLV-1 WT PBS (E and G) or Δ425-434 PBS (F and H) in the presence of 2 μM HTLV-1 WT Gag (E and F) or ΔC29 Gag (G and H) at 37°C. (I and J): Graphs showing percentages of tRNA^Pro^ annealed to HTLV-1 WT PBS (I) or Δ425-434 PBS (J) at different time points in the presence of 2 μM HTLV-1 WT or ΔC29 Gag. Lines represent exponential fits of the data with the standard deviation between trials indicated.

**Table 1.**
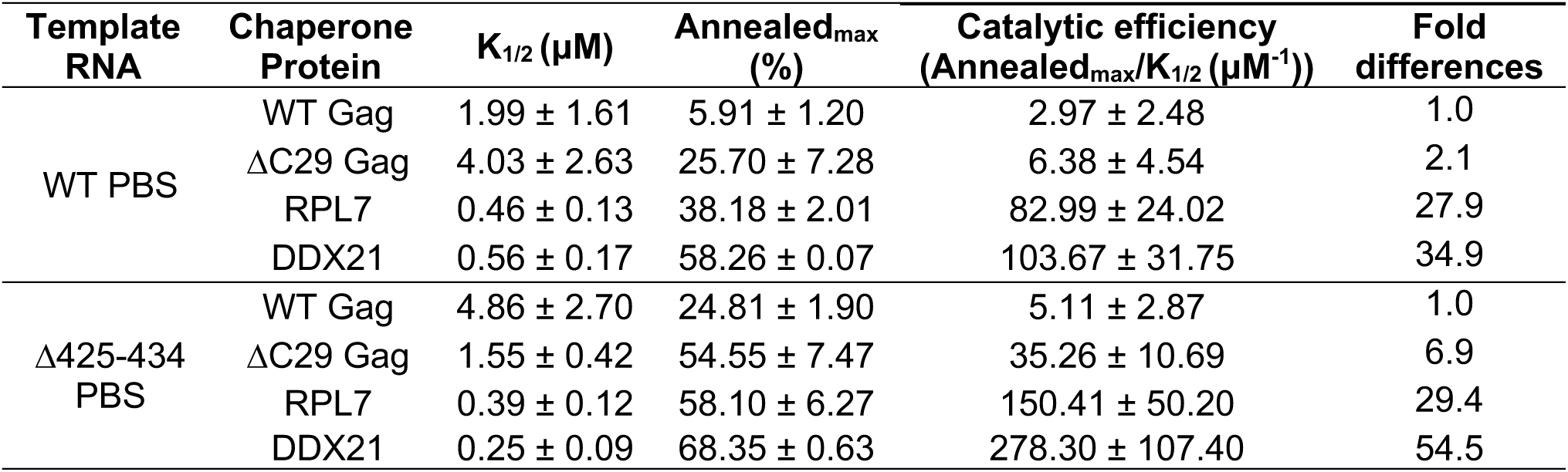
Summary of K_1/2_ (μM), Annealed_max_ (%), annealing efficiencies (Annealed_max_/K_1/2_ (μM^-1^)), and fold differences of catalytic efficiencies of chaperone proteins tested in concentration-dependence annealing assays.

| Template RNA | Chaperone Protein | $K_{1/2}$ ($\mu\text{M}$) | $\text{Annealed}_{\text{max}}$ (%) | Catalytic efficiency ($\text{Annealed}_{\text{max}}/K_{1/2}$ ($\mu\text{M}^{-1}$)) | Fold differences |
| --- | --- | --- | --- | --- | --- |
| WT PBS | WT Gag | $1.99 \pm 1.61$ | $5.91 \pm 1.20$ | $2.97 \pm 2.48$ | 1.0 |
| | $\Delta\text{C29}$ Gag | $4.03 \pm 2.63$ | $25.70 \pm 7.28$ | $6.38 \pm 4.54$ | 2.1 |
| | RPL7 | $0.46 \pm 0.13$ | $38.18 \pm 2.01$ | $82.99 \pm 24.02$ | 27.9 |
| | DDX21 | $0.56 \pm 0.17$ | $58.26 \pm 0.07$ | $103.67 \pm 31.75$ | 34.9 |
| $\Delta\text{425-434}$ PBS | WT Gag | $4.86 \pm 2.70$ | $24.81 \pm 1.90$ | $5.11 \pm 2.87$ | 1.0 |
| | $\Delta\text{C29}$ Gag | $1.55 \pm 0.42$ | $54.55 \pm 7.47$ | $35.26 \pm 10.69$ | 6.9 |
| | RPL7 | $0.39 \pm 0.12$ | $58.10 \pm 6.27$ | $150.41 \pm 50.20$ | 29.4 |
| | DDX21 | $0.25 \pm 0.09$ | $68.35 \pm 0.63$ | $278.30 \pm 107.40$ | 54.5 |

To test the effect of the PBS context on annealing inhibition, we deleted the sequence (nt 425-434) on one side of the stem in the PBS hairpin (Figure 1B, Δ425-434 PBS). We hypothesized that deleting this sequence would facilitate Gag-chaperoned annealing by making the PBS region less structured and more accessible. In concentration-dependence tRNA^Pro^ annealing assays with the Δ425-434 PBS and HTLV-1 WT or ΔC29 Gag, significantly more complex formation was observed compared to the WT PBS (Figure 2A – 2D). This trend was also observed in the heat-annealed positive control. Band intensity for the B1 complex was more prominent for tRNA^Pro^ heat-annealed to the Δ425-434 PBS (Figure 2B) than to the WT PBS (Figure 2A). These results confirmed our hypothesis that the Δ425-434 PBS is more accessible resulting in greater tRNA^Pro^ annealing. As summarized in Table 1, in the presence of 4 μM HTLV-1 WT Gag, 24.8 ± 1.9% of tRNA^Pro^ is annealed to the Δ425-434 PBS, K_1/2_ is 4.86 ± 2.7 μM, and the catalytic efficiency is 5.11 ± 2.9 μM^-1^ (Figure 2B and 2D; Table 1). In contrast, with 4 μM HTLV-1 ΔC29 Gag, 54.6 ± 7.5% of tRNA^Pro^ is annealed to the Δ425-434 PBS, K_1/2_ is 1.55 ± 0.42 μM, and the catalytic efficiency is 35.3 ± 11 μM^-1^ (Figure 2B and 2D; Table 1). HTLV-1 ΔC29 Gag chaperoned tRNA^Pro^ annealing to the Δ425-434 PBS 6.9-fold more efficiently than WT Gag (Table 1).

Time-course annealing assays were also performed in the presence of 2 μM HTLV-1 WT or ΔC29 Gag (Figure 2E – 2J). After a 1-h incubation with the WT PBS, 1.52 ± 0.19% of tRNA^Pro^ is annealed by WT Gag, whereas 31.0 ± 2.4% is annealed by ΔC29 Gag (Figure 2E, 2G, 2I; Table 2). The annealing rate (k) for ΔC29 Gag is 0.060 ± 0.008 min^-1^. The scaled annealing rate (k’), which was calculated by multiplying k by the fraction of tRNA^Pro^ annealed, is 0.019 ± 0.003 min^-1^ (Figure 2G and 2I; Table 2). The annealing rate for WT Gag could not be determined due to its low efficiency (Figure 2E). For the Δ425-434 PBS time-course annealing assays, 18.4 ± 9.7% of tRNA^Pro^ is annealed by WT Gag, whereas 54.9 ± 8.3% is annealed by ΔC29 Gag (Figure 2F, 2H, 2J; Table 2). The scaled k’ values for WT and ΔC29 Gag are 0.022 ± 0.02 min^-1^ and 0.136 ± 0.028 min^-1^, respectively (Figure 2F, 2H, 2J; Table 2). The trend for time-course annealing assays is similar to concentration-dependence annealing assays. Overall, HTLV-1 WT or ΔC29 Gag facilitated tRNA^Pro^ annealing to the Δ425-434 PBS more efficiently than to the WT PBS. For the Δ425-434 PBS, ΔC29 Gag catalyzed tRNA^Pro^ annealing 6.2-fold faster than WT Gag (Table 2).

**Table 2.**
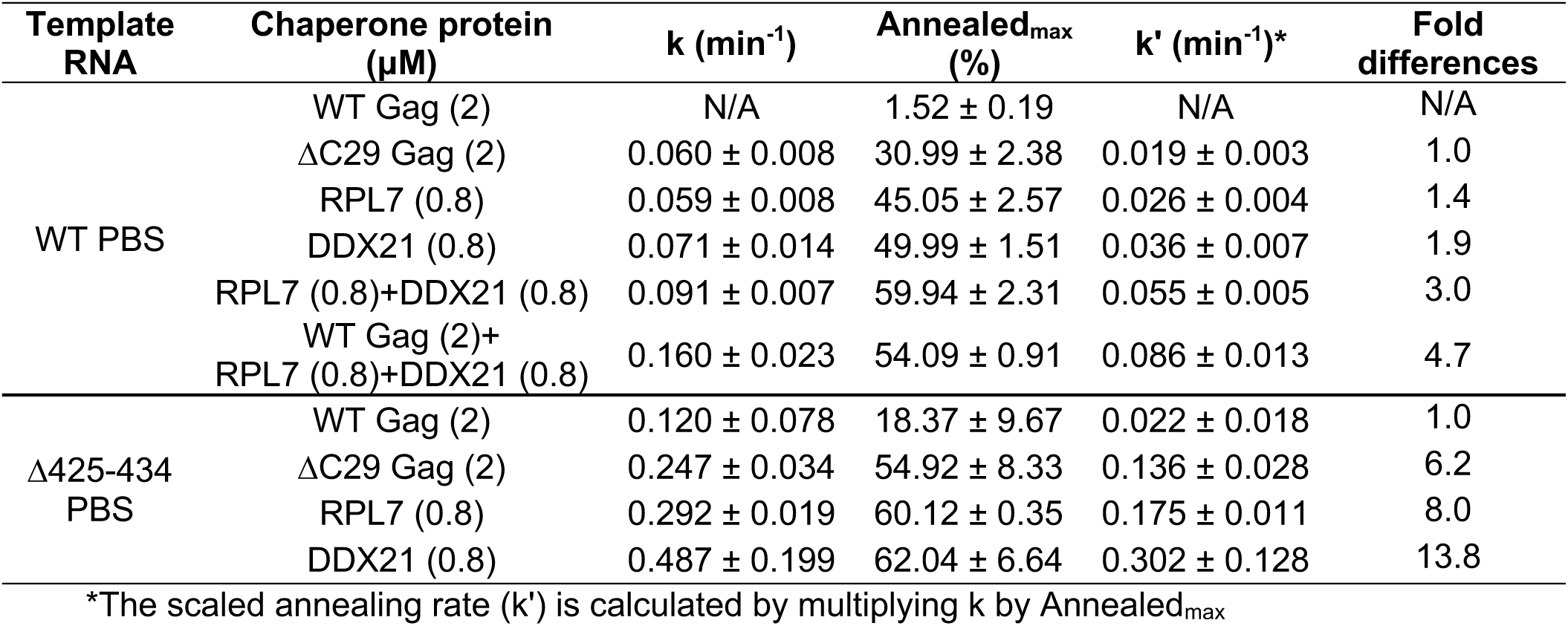
Summary of annealing rates (k (min^-1^)), Annealed_max_ (%), scaled annealing rates (k’ (min^-1^)), and fold differences of scaled annealing rates of chaperone proteins tested in time-course annealing assays.

| Template RNA | Chaperone protein ($\mu\text{M}$) | $k$ ($\text{min}^{-1}$) | Annealed<sub>max</sub> (%) | $k'$ ($\text{min}^{-1}$)* | Fold differences |
| --- | --- | --- | --- | --- | --- |
| WT PBS | WT Gag (2) | N/A | $1.52 \pm 0.19$ | N/A | N/A |
| | $\Delta\text{C29}$ Gag (2) | $0.060 \pm 0.008$ | $30.99 \pm 2.38$ | $0.019 \pm 0.003$ | 1.0 |
| | RPL7 (0.8) | $0.059 \pm 0.008$ | $45.05 \pm 2.57$ | $0.026 \pm 0.004$ | 1.4 |
| | DDX21 (0.8) | $0.071 \pm 0.014$ | $49.99 \pm 1.51$ | $0.036 \pm 0.007$ | 1.9 |
| | RPL7 (0.8)+DDX21 (0.8) | $0.091 \pm 0.007$ | $59.94 \pm 2.31$ | $0.055 \pm 0.005$ | 3.0 |
| | WT Gag (2)+<br>RPL7 (0.8)+DDX21 (0.8) | $0.160 \pm 0.023$ | $54.09 \pm 0.91$ | $0.086 \pm 0.013$ | 4.7 |
| $\Delta\text{425-434}$<br>PBS | WT Gag (2) | $0.120 \pm 0.078$ | $18.37 \pm 9.67$ | $0.022 \pm 0.018$ | 1.0 |
| | $\Delta\text{C29}$ Gag (2) | $0.247 \pm 0.034$ | $54.92 \pm 8.33$ | $0.136 \pm 0.028$ | 6.2 |
| | RPL7 (0.8) | $0.292 \pm 0.019$ | $60.12 \pm 0.35$ | $0.175 \pm 0.011$ | 8.0 |
| | DDX21 (0.8) | $0.487 \pm 0.199$ | $62.04 \pm 6.64$ | $0.302 \pm 0.128$ | 13.8 |

\*The scaled annealing rate ( $k'$ ) is calculated by multiplying $k$ by Annealed<sub>max</sub>

### Affinity tagging/purification-mass spectrometry (AP-MS) identifed HTLV-1 Gag interacting partners

Although HTLV-1 ΔC29 Gag chaperoned tRNA^Pro^ annealing to the PBS more efficiently than WT Gag, this Gag truncation variant is not known to be present in HTLV-1 virions. In fact, the acidic C-terminus of HTLV-1 NC prevents APOBEC3G from being packaged into virions and thus counteracts the APOBEC3G restriction (31). We hypothesize that host co-factors of HTLV-1 Gag may help increase its chaperone activity. To identify HTLV-1 Gag interacting partners, an AP-MS-based proteomics analysis was performed. C-terminal GFP-tagged HTLV-1 Gag or an empty GFP expression vector (as a pull-down background control) was expressed in HEK293T cells. This vector encodes a twin-strep tag at the C-terminus of GFP, which is pulled down by streptactin (Figure 1D). Using this strategy, both GFP and HTLV-1 Gag-GFP were significantly enriched in the pull-down fractions (Figure S2A) and a large number of co-factors were identified by MS.

A list of all hits can be found in Table S3 and the top 10 protein hits are listed in Table 3. Among the top hits, six are ribosomal proteins (RPL4, RPL7A, RPL6, RPL7, RPS3A, and RPL18), two are protein glycosyltransferases (RPN1 and DDOST), two are NA chaperone proteins/helicases (RPL7 and DDX21), and one is fatty acid peroxisomal importer protein (ABCD3). Both RPN1 and DDOST are subunits in the oligosaccharyltransferase (OST) complex, which is located in the rough ER and essential for N-linked glycosylation on asparagine residues (32). Ribosomal protein L7 (RPL7) has been reported to be packaged into HIV-1 virions and to interact with the NC domain of HIV-1 Gag to synergistically increase annealing of tRNA^“#$%^to the PBS (21,22). Nucleolar RNA helicase 2 (DDX21) is an ATP-dependent helicase and displays helix-unwinding and RNA-folding activities (33,34). It was shown to interact with HIV-1 Rev through the DEAD domain and thereby increase Rev-RRE binding affinity (35). Because our goal was to identify potential RNA chaperone proteins and/or helicases that may help increase HTLV-1 Gag’s chaperone activity, we picked two known chaperones/helicases identified in the AP-MS study for validation and investigation as potential tRNA annealing factors: RPL7 and DDX21.

**Table 3.**
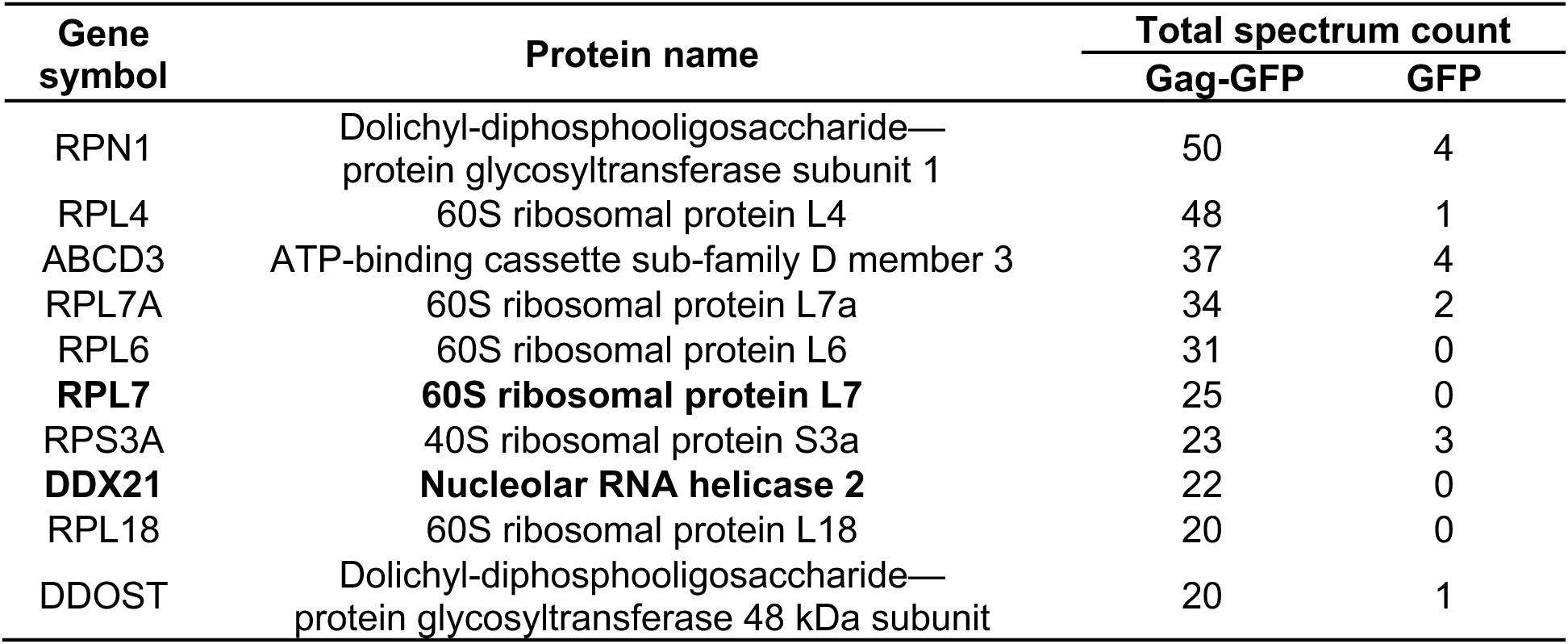
List of top 10 AP-MS protein hits.

### Interactions of HTLV-1 Gag with RPL7 and DDX21 are RNA-independent and confirmed by pull-down and reciprocal co-immunoprecipitation (co-IP)

To confirm the AP-MS results, HTLV-1 Gag-GFP pull-down and HTLV-1 Gag-FLAG co-IP followed by immunoblotting were performed in HEK293T cell lysate. When HTLV-1 Gag-GFP, but not GFP, was pulled-down, RPL7 and DDX21 were co-pulled-down (Figure 3A). When HTLV-1 Gag-FLAG, but not FLAG, was IP’d, RPL7 and DDX21 were co-IP’d (Figure 3B). Both RPL7 and DDX21 are RNA-binding proteins (35,36). To investigate whether the interactions of HTLV-1 Gag with RPL7 and DDX21 are direct or mediated by RNA, cell lysate was treated with RNase A/T1 before Gag-GFP pull-down assays. We found that the levels of RPL7 and DDX21 co-pulled-down with Gag-GFP were not impacted by RNase treatment suggesting the interactions are RNA-independent (Figure 3A). To further confirm the interaction, a reciprocal co-IP was performed. In cell lysate from HEK293T cells co-overexpressing Gag-GFP and HA-RPL7 (Figure 1D and 1F), when HA-RPL7 was IP’d, Gag-GFP and DDX21 were co-IP’d. No interaction was observed in the IgG isotype background control sample (Figure 3D). To rule out the possibility that RPL7 interacts with DDX21 through Gag, HA-RPL7 co-IP was performed in HEK293T cell lysate without Gag expression. DDX21 was co-IP’d with HA-RPL7 in the absence of Gag (Figure 3E). Another reciprocal DDX21 co-IP in HEK293T cells overexpressing Gag-GFP was performed, DDX21 IP’d by anti-DDX21 antibody was co-IP’d with Gag-GFP and RPL7 (Figure 3F). The above results were obtained from exogenously overexpressed Gag in HEK293T cells. To confirm the interactions at the endogenous expression levels, co-IP was carried out in a more physiologically relevant chronically HTLV-1-infected MT-2 cell line. When Gag, CA-NC, and CA were IP’d by an anti-p24 antibody, RPL7 and DDX21 were co-IP’d (Figure 3C). Overall, these results confirm the interactions of HTLV-1 Gag with DDX21 and RPL7 and are consistent with the AP-MS data. In addition, RPL7 and DDX21 also interact with each other, indicating that Gag, RPL7, and DDX21 may form a complex.

**Figure 3.**
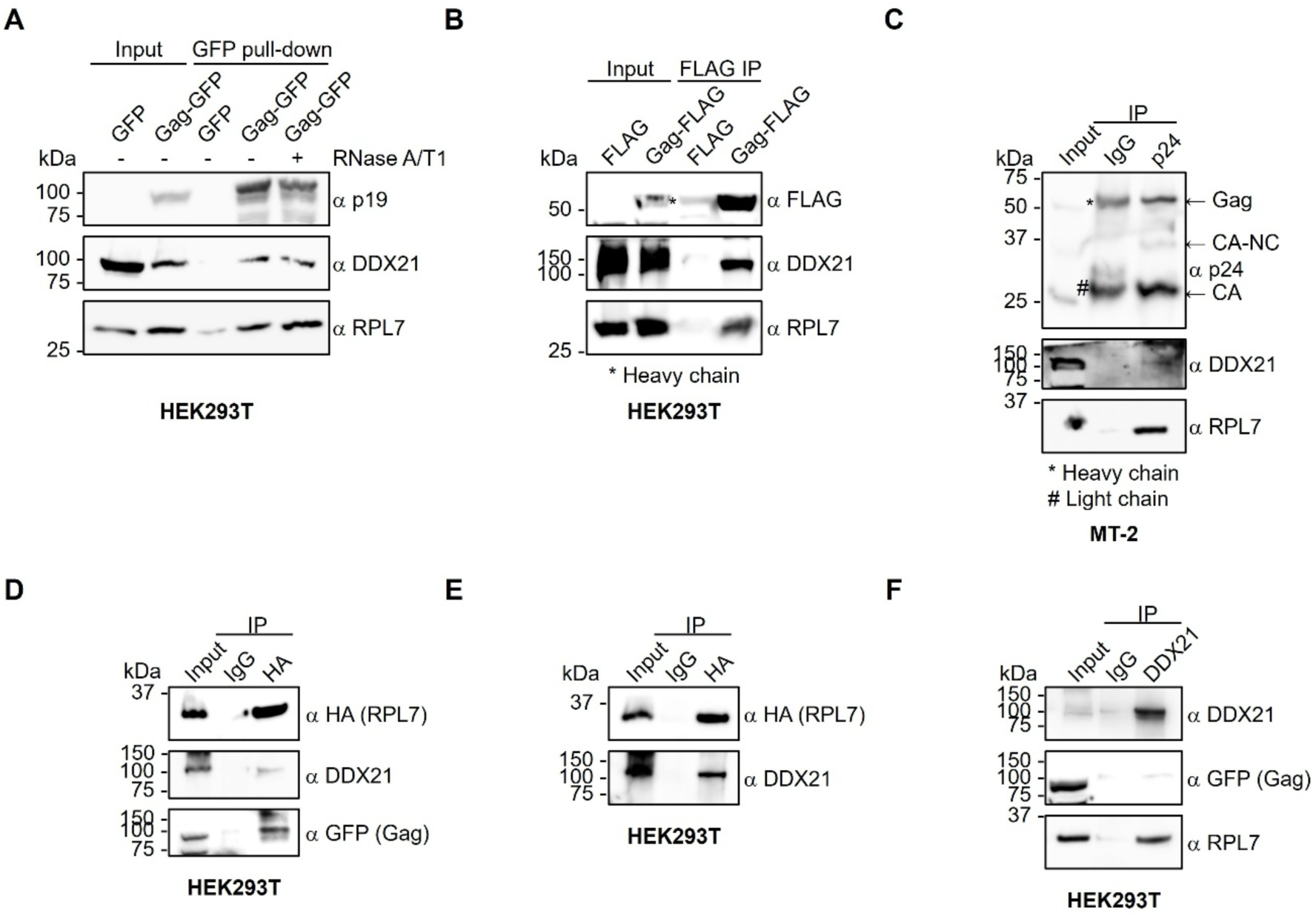
HTLV-1 Gag interacts with RPL7 and DDX21 in an RNA-independent manner as confirmed by pull-down and reciprocal co-IP. (A) Western blot analysis of HEK293T cell lysates overexpressing GFP or HTLV-1 Gag-GFP with or without RNase treatment. Proteins were pulled down by streptactin and immunoblotted using anti-HTLV-1 p19 (MA), anti-DDX21 and anti-RPL7 antibodies. (B) FLAG or HTLV-1 Gag-FLAG expressed in HEK293T cell lysate was IP’d using an anti-FLAG antibody. Co-IP’d proteins were immunoblotted using anti-DDX21 and anti-RPL7 antibodies. (C) Gag, CA-NC, and CA in MT-2 cell lysate were IP’d using an anti-HTLV-1 p24 (CA) antibody. Co-IP’d proteins were immunoblotted using anti-DDX21 and anti-RPL7 antibodies. (D and E): RPL7-HA was IP’d using an anti-HA antibody in lysate from HEK293T cells with (D) or without (E) HTLV-1 Gag-GFP co-overexpression. Co-pulled-down proteins were immunoblotted using anti-DDX21 and anti-GFP antibodies. (F) DDX21 was IP’d by an anti-DDX21 antibody in lysate from HEK293T cells overexpressing HTLV-1 Gag-GFP. Co-immunoprecipitated proteins were immunoblotted using anti-GFP and anti-RPL7 antibodies.

HTLV-1 Gag is essential for assembling viral particles and packaging gRNA into virions. In addition to specifically binding to gRNA, two other features of Gag, myristoylation and oligomerization, are required for efficient gRNA packaging. Post-translational myristoylation of the second residue, Gly2, in the MA domain of Gag is required for Gag trafficking from the cytoplasm to the PM and anchoring to the PM (28,37). Gag oligomerization occurs on or near the PM through N-terminal CA domain interactions (38). The residues reported to be involved in oligomerization are Met147 and Tyr191 located in the CA dimer interface, and Gln177 and Phe178 located in the CA trimer interface (39). To test the role of myristoylation, Gly2 was changed to Ala (G2A). To disrupt Gag oligomer formation, M147AY191A and Q177AF178A mutants were made. We observed that the levels of RPL7 and DDX21 co-IP’d with Gag-FLAG were not changed significantly in these myristoylation- and oligomerization-deficient Gag mutants in comparison to WT Gag (Figure S3A). These results demonstrate that interactions of HTLV-1 Gag with RPL7 and DDX21 are direct and independent of the presence of RNA, Gag membrane trafficking, and Gag multimerization.

### The NC zinc finger (ZF) structures of HTLV-1 Gag are essential for the interactions with RPL7 and DDX21

Next, we investigated which domain and subdomain in HTLV-1 Gag interact with RPL7 and DDX21. Single-domain truncation mutants of Gag-GFP (ΔMA, ΔCA, and ΔNC) were prepared and transfected into HEK293T cells, followed by Gag-GFP pull-down and immunoblotting (Figure 4A). The interactions of Gag with RPL7 and DDX21 were significantly diminished only upon deletion of the NC domain (Figure 4B). The NC domain mainly consists of three subdomains, zinc finger 1 (ZF1), zinc finger 2 (ZF2), and the highly acidic C-terminal domain (CTD) (Figure 4A). To further map which subdomains interact with RPL7 and DDX21, a series of single-or double-subdomain truncation mutants, ΔZF1, ΔZF2, ΔZF1-2, and ΔNC CTD were made and transfected into HEK293T cells, followed by pull-down assays (Figure 4A). In comparison to the interactions of WT Gag with RPL7 and DDX21, the interaction was slightly reduced when ZF1 in Gag was deleted and was more severely impaired when ZF2 or both ZF1 and ZF2 were absent (Figure 4C). These results indicate that ZF2 is more important than ZF1 for the interaction and that HTLV-1 Gag interacts with RPL7 and DDX21 through both ZFs.

**Figure 4.**
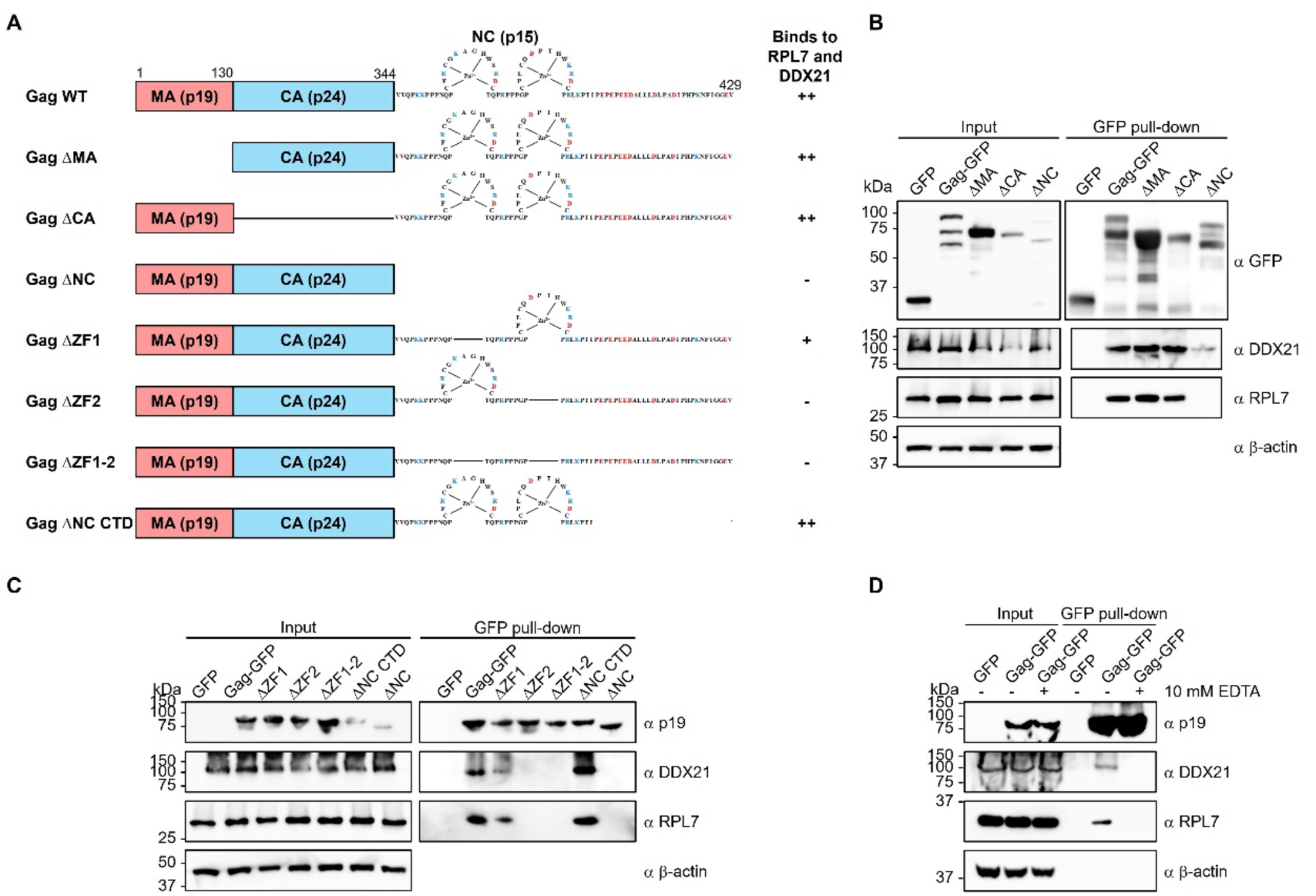
Zinc finger (ZF) structures in the nucleocapsid (NC) domain of HTLV-1 Gag are essential for the interactions with DDX21 and RPL7. (A) Schematic representation of HTLV-1 Gag and NC subdomain truncation constructs used in the pull-down study. (++), (+), and (-) indicate the relative ability of each construct to co-pull-down DDX21 and RPL7. (B) GFP, WT Gag-GFP, ΔMA, ΔCA, and ΔNC Gag-GFP in HEK293T cell lysate were pulled down by streptactin and immunoblotted using an anti-GFP antibody 48 h post-transfection. Co-pulled-down proteins were immunoblotted using anti-DDX21 and anti-RPL7 antibodies. (C) GFP, WT Gag-GFP, ΔZF1, ΔZF2, ΔZF1-2, ΔNC CTD, and ΔNC Gag-GFP in HEK293T cell lysate were pulled down by streptactin and immunoblotted using an anti-HTLV-1 p19 antibody 48 h post-transfection. Co-pulled-down proteins were immunoblotted using anti-DDX21 and anti-RPL7 antibodies. (D) Cell lysate from HEK293T cells overexpressing HTLV-1 Gag-GFP 48 h post-transfection was treated with or without 10 mM EDTA. GFP or HTLV-1 Gag-GFP was pulled down by streptactin and immunoblotted using an anti-HTLV-1 p19 antibody. Co-pulled-down proteins were immunoblotted using anti-DDX21 and anti-RPL7 antibodies. β-actin was used as a loading control in panels (B - D).

The coordination of Zn^2+^ is crucial for maintaining NC’s structure (40). To investigate the importance of ZF structures on the interaction, 10 mM EDTA was added to the cell lysate as a Zn^2+^ chelator prior to performing Gag-GFP pull-down assays (Figure 4D). The interactions of Gag-GFP with RPL7 and DDX21 were abolished upon EDTA treatment suggesting that the ZF structures in the NC domain of HTLV-1 Gag are indispensable for interacting with RPL7 and DDX21.

### N-terminal basic leucine zipper (bZIP) domain and C-terminal domain (CTD) of RPL7 interact with HTLV-1 Gag and DDX21

We next probed which domains in RPL7 are involved in HTLV-1 Gag and DDX21 interactions. RPL7 is composed of the N-terminal basic leucine zipper domain (bZIP, residues 1-77), the C-terminal nucleic acid-binding domain (CTD, residues 198-248), and the central globular domain (residues 78-197) (Figure 5A) (36,41). The CTD lacks a canonical NA-binding domain, and bZIP and CTD have different preferences for RNA binding (36,41,42). The bZIP domain has a higher binding affinity to mRNA than 28S rRNA (41), whereas the opposite is true for the CTD (36). Other known RPL7 cellular interacting partners include ribosomal protein S7 (RPS7), ZFN7, and vitamin D receptor (VDR). RPL7 interacts with all of them through the bZIP domain (43,44). To map which domains in RPL7 interact with HTLV-1 Gag or DDX21, a series of single-domain (Figure 5A, constructs B to D) or double-domain (Figure 5A, constructs E to G) truncation mutants were prepared. In mutant C, the (GGGGS)_3_ flexible linker was added between the N-terminal bZIP and CTD to separate these two domains and minimize interfering with domain folding (Figure 5A, construct C) (45). Mutant G contains the coding sequence for only the C-terminal 51 residues; it failed to express in HEK293T cells, so we were unable to assess its influence on the interactions (Figure 5B and 5C).

**Figure 5.**
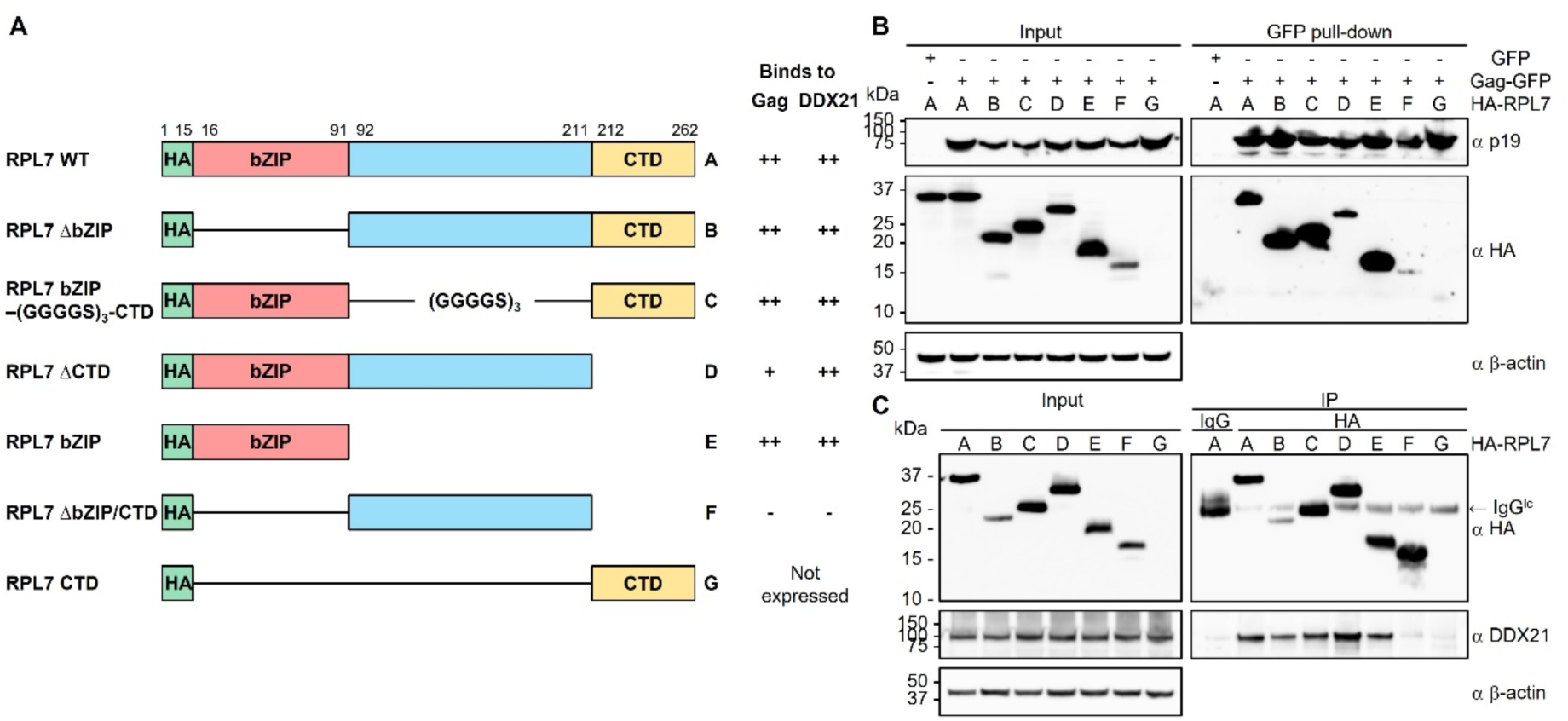
RPL7 interacts with HTLV-1 Gag and DDX21 through the N-terminal basic leucine zipper (b-ZIP) domain and C-terminal domain (CTD). (A) Schematic representation of human RPL7 WT and domain truncation constructs used in this study. On the right side, letters indicate the construct used in each lane in (B) and (C). Relative binding of RPL7 variants to HTLV-1 Gag and DDX21 are shown as (++), (+), or (-). (B) GFP or Gag-GFP was pulled down by streptactin and immunoblotted using an anti-HTLV-1 p19 antibody in cell lysate from HEK293T cells co-transfected with HTLV-1 Gag-GFP and WT or truncation mutant constructs of HA-RPL7 48 h post-transfection. Co-pulled-down HA-RPL7 was immunoblotted using an anti-HA antibody. (C) RPL7-HA WT or truncation mutants were IP’d and immunoblotted using anti-HA antibody 48 h post-transfection. Co-IP’d DDX21 was immunoblotted using anti-DDX21 antibody. (B and C): β-actin was used as a loading control.

To study which domains in RPL7 interact with HTLV-1 Gag, Gag-GFP and WT or truncation mutant constructs of HA-RPL7 were co-transfected into HEK293T cells and Gag-GFP pull-down assays were performed (Figure 5B). In comparison to the amount of WT HA-RPL7 co-pulled-down with Gag-GFP, the interaction was slightly decreased in mutant D lacking the CTD and severely impaired in mutant F when both NA-binding domains were deleted (Figure 5B). These results show that CTD plays a more crucial role than bZIP for the interaction. However, bZIP also participates in the interaction as a construct lacking both bZIP and CTD results in a more severely impaired interaction compared with the single-domain CTD deletion. To examine which domains in RPL7 interact with DDX21, WT or truncation mutant constructs of HA-RPL7 were transfected into HEK293T cells and HA-RPL7 co-IP was carried out. Similar to results obtained with the HTLV-1 Gag/HA-RPL7 study, the interaction between DDX21 and RPL7 was severely impaired with the ΔbZIP/CTD RPL7 mutant (Figure 5C, construct F). Taken together, both NA-binding domains, bZIP and CTD, in RPL7 interact with HTLV-1 Gag and DDX21.

### Core helicase and C-terminal domains of DDX21 interact with HTLV-1 Gag

DDX21 is a member of the DEAD-box helicase family. The domain organization of DDX21 includes a disordered N-terminal domain (NTD, residues 1-116), an ATP-binding DEAD-box helicase N-terminal domain (helicase N, residues 117-341), a helicase C-terminal domain (helicase C, residues 342-498), and a disordered C-terminal foldase domain (CTD, residues 499-715) (Figure 6A) (33). DDX21 displays two distinct enzymatic activities; its ATP-dependent RNA-unwinding activity requires the disordered N-terminus and the core helicase domain, while its RNA folding activity requires the C-terminal foldase domain. (33,34). To map which domains in DDX21 interact with HTLV-1 Gag, a series of single-domain (Figure 6A, constructs B to E) and double-domain (Figure 6A, constructs F and G) deletion DDX21 mutants were prepared. HTLV-1 Gag-GFP and WT or deletion mutants of mCherry-DDX21-V5 were co-transfected into HEK293T cells and analyzed in GFP-Gag pull-down studies. In comparison to the amount of WT DDX21 co-pulled-down with Gag-GFP, the interaction was slightly decreased in mutant D lacking the helicase C-terminal domain and completely abolished in mutant F when the helicase core, containing both helicase N- and C-terminal domains, were missing (Figure 6B). These results indicate that the presence of both domains in the helicase core is essential for the interaction, and that the helicase C-terminal domain is more important for interacting with Gag than the helicase N-terminal domain. In addition, the interaction disappeared when the CTD was absent in mutant E (Figure 6B). Collectively, DDX21 interacts with HTLV-1 Gag through the helicase core and the CTD.

**Figure 6.**
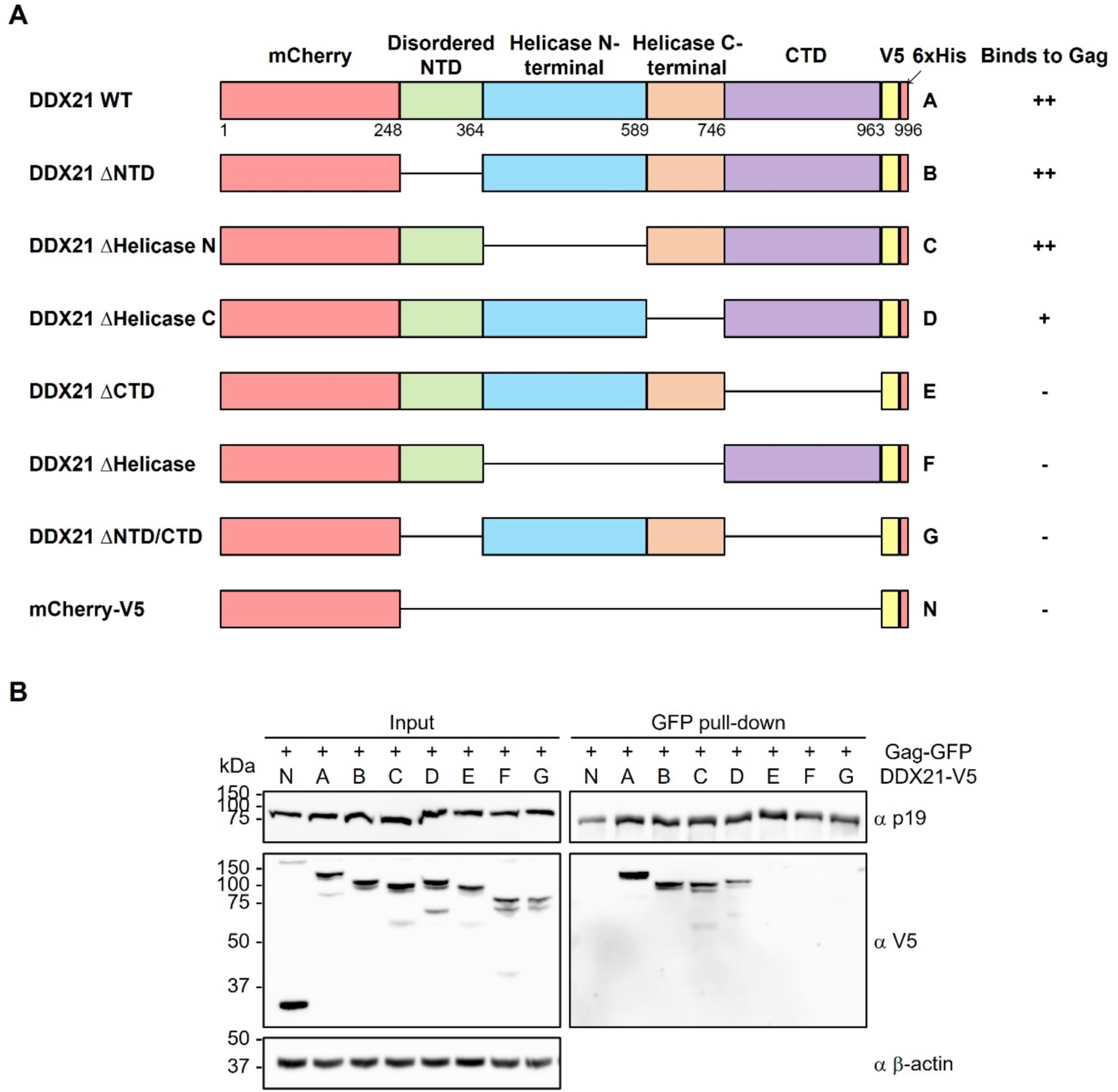
DDX21 interacts with HTLV-1 Gag through the helicase core and the C-terminal domain. (A) Schematic representation of human DDX21 WT and domain truncation constructs used in the pull-down study. Letters on the right represent the construct used in each lane in (B). Relative binding of DDX21 variants to HTLV-1 Gag are reported as (++), (+), or (-). (B) Gag-GFP was pulled down by streptactin and immunoblotted using an anti-HTLV-1 p19 antibody in cell lysate from HEK293T cells co-transfected with HTLV-1 Gag-GFP and WT or truncation mutant constructs of mCherry-DDX21-V5 48 h post-transfection. Co-pulled-down DDX21-V5 was immunoblotted using an anti-V5 antibody. β-actin was used as a loading control.

### RPL7 and DDX21 are packaged into HTLV-1 virions

To investigate whether these Gag-interacting partners are packaged into virions, virions purified by sucrose cushion were further fractionated using an OptiPrep density gradient to remove secreted proteins, exosomes, and cytoplasmic contaminants (Figure 7A) (46). As shown in the top blot probed with anti-HTLV-1 p24 antibody in Figure 7B, HTLV-1 virions were present in fractions 9 to 5 following ultracentrifugation. Both RPL7 and DDX21 co-sedimented with CA and Gag in these same fractions. The ratio of RPL7 or DDX21 to Gag was highest in the larger virus particles (Figure 7B, fraction 6).

**Figure 7.**
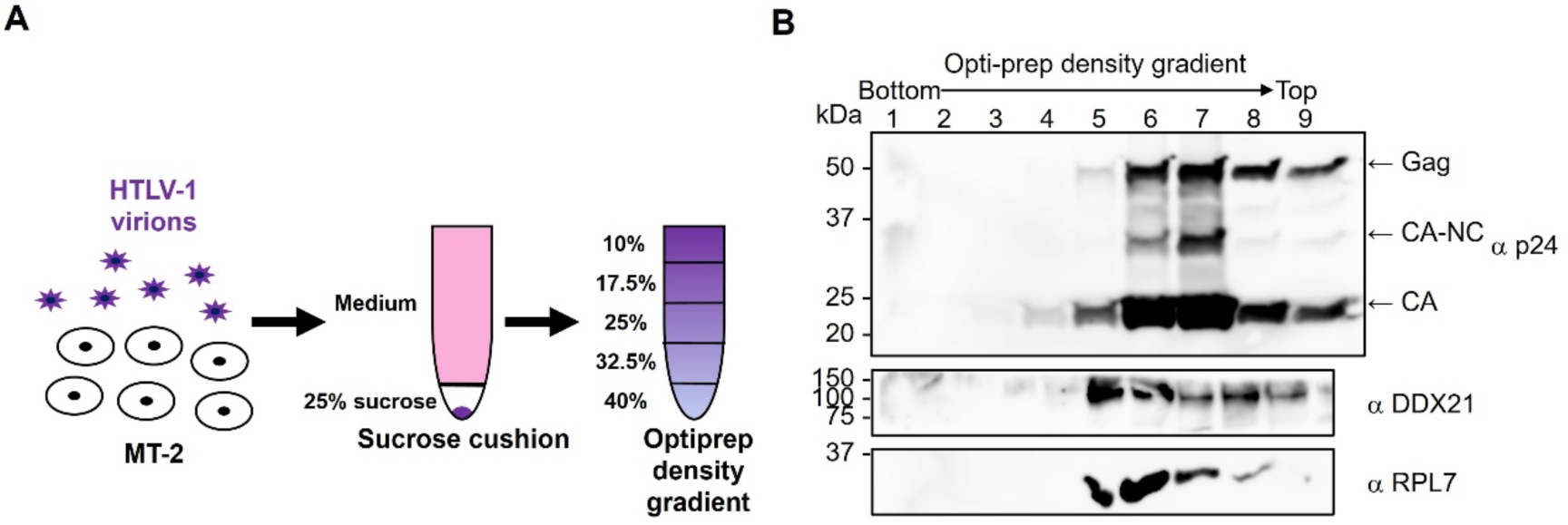
RPL7 and DDX21 are packaged into HTLV-1 virions. (A) Schematic diagram of purification of HTLV-1 virus from MT-2 cell culture medium through a sucrose cushion and further fractionation of the viral pellet through an OptiPrep density gradient. (B) Fractionated viral lysate was immunoblotted using anti-HTLV-1 p24, anti-DDX21, and anti-RPL7 antibodies.

### Human DDX21 and RPL7 facilitate tRNA^Pro^ annealing to the HTLV-1 PBS more efficiently than HTLV-1 WT and ΔC29 Gag

We next purified RPL7 and DDX21 proteins and characterized their ability to promote tRNA^Pro^ annealing to the HTLV-1 PBS. WT and mutant RPL7 lacking one or both NA-binding domains, and WT DDX21 proteins were purified from *E. coli* (Figure S4A and S4B). To confirm that purified WT RPL7 protein was folded properly, circular dichroism (CD) was performed. The spectrum shows two minima at 208 nm and 222 nm corresponding to the characteristic signals for α-helical proteins (Figure S4C) (47). This result is consistent with the α-helix-rich RPL7 structure solved by cryo-electron microscopy (cryo-EM) (48).

To characterize the chaperone activities of RPL7 and DDX21, tRNA^Pro^ annealing assays were performed with the same RNA constructs shown in Figure 1A. In concentration-dependence annealing assays carried out for a period of 60 min (Figure 8A -8D), when more RPL7 or DDX21 proteins were present in the annealing reactions, less free tRNA^Pro^ was detected and more B1 and B2 complexes formed. RPL7 and DDX21 facilitate tRNA^Pro^ annealing to the PBS more efficiently than HTLV-1 WT or ΔC29 Gag (compare Figures 2A and 2B with Figures 8A - 8D). With 4 μM chaperone protein and the WT PBS in the reaction, RPL7 promotes 38 ± 2% of tRNA^Pro^ annealed while DDX21 achieves 58 ± 0.07% complex formation (Figure 8A and 8C). K_1/2_ for RPL7 is 0.46 ± 0.13 μM and the catalytic efficiency (Annealed_max_/K_1/2_) is 83 ± 24 μM^-1^ (Table 1). For DDX21, K_1/2_ is 0.56 ± 0.17 μM and the catalytic efficiency is 104 ± 32 μM^-1^ (Table 1). With 4 μM chaperone protein and the less structured Δ425-434 PBS in the reaction, RPL7 promotes 58 ± 6% of tRNA^Pro^ annealing whereas 68 ± 0.6% of tRNA^Pro^ is annealed by DDX21 (Figure 8B and 8D). K_1/2_ for RPL7 is 0.39 ± 0.12 μM and the catalytic efficiency is 150 ± 50 μM^-1^ (Table 1). For DDX21, K_1/2_ is 0.25 ± 0.09 μM and the catalytic efficiency is 278 ± 107 μM^-1^ (Table 1). In summary, DDX21 chaperoned tRNA^Pro^ annealing 1.3- and 16-fold more effectively to the WT PBS and 1.9- and 7.9-fold more effectively to the Δ425-434 PBS compared to RPL7 and ΔC29 Gag, respectively. RPL7 facilitated tRNA^Pro^ annealing 13-fold more efficiently to the WT PBS and 4.3-fold more efficiently to the Δ425-434 PBS compared to ΔC29 Gag (Table 1).

**Figure 8.**
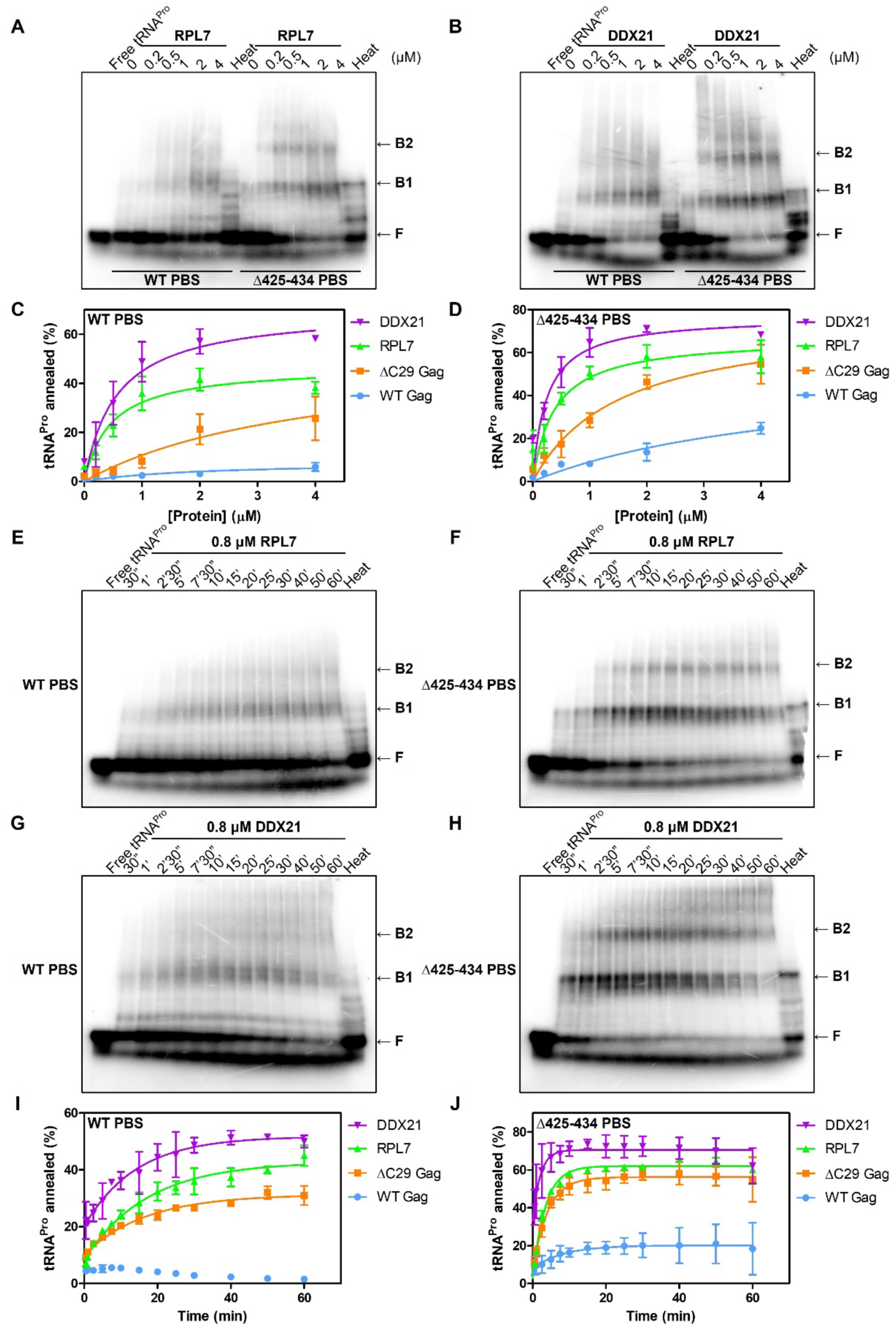
Concentration-dependence and time-course annealing assays show that human DDX21 and RPL7 facilitate tRNA^Pro^ annealing to the HTLV-1 PBS more efficiently than HTLV-1 WT and ΔC29 Gag. (A and B): Concentration-dependence annealing assays using 20 nM 5’ ^32^P-labeled tRNA^Pro^ and 200 nM HTLV-1 WT PBS or Δ425-434 PBS in the presence of varying concentrations of RPL7 (A) or DDX21 (B) at 37°C for 1 h. F indicates free tRNA^Pro^ and B1 and B2 indicate different tRNA^Pro^-PBS binary complexes. Heat indicates heat annealing, the positive control. (C and D): Graphs showing percentages of tRNA^Pro^ annealed to WT (C) or Δ425-434 PBS (D) in the presence of varying concentrations of HTLV-1 WT Gag, ΔC29 Gag, RPL7, or DDX21. (E-H): Time-course annealing assays using 20 nM 5’ ^32^P-labeled tRNA^Pro^ and 200 nM HTLV-1 WT PBS (E and G) or Δ425-434 PBS (F and H) in the presence of 0.8 μM RPL7 (E and F) or DDX21 (G and H) at 37°C. (I and J): Graphs showing percentages of tRNA^Pro^ annealed to HTLV-1 WT PBS (I) or Δ425-434 PBS (J) at different time points in the presence of 2 μM HTLV-1 WT Gag, 2 μM ΔC29 Gag, 0.8 μM RPL7, or 0.8 μM DDX21. Lines represent exponential fits of the data with the standard deviation between trials indicated.

Time-course annealing assays were next performed in the presence of 0.8 μM RPL7 or DDX21; reactions were terminated at different time points up to 1 h (Figure 8E - 8H). After a 1-h incubation with the WT PBS, 45 ± 3% of tRNA^Pro^ is annealed by RPL7, while 50 ± 2% of tRNA^Pro^ is annealed by DDX21. The scaled annealing rates (k’) are 0.026 ± 0.004 min^-1^ and 0.036 ± 0.007 min^-1^, respectively (Figure 8E, 8G, 8I; Table 2). For the Δ425-434 PBS time-course annealing assays, 60 ± 0.4% of tRNA^Pro^ is annealed by RPL7, while 62 ± 7% of tRNA^Pro^ is annealed by DDX21. The k’ values for RPL7 and DDX21 are 0.18 ± 0.01 min^-1^ and 0.30 ± 0.13 min^-1^ (Figure 8F, 8H, 8J; Table 2). The trend for time-course annealing assays is similar to concentration-dependence assays; 0.8 μM RPL7 and DDX21 chaperoned tRNA^Pro^ annealing to the Δ425-434 PBS more efficiently than to the WT PBS. Overall, DDX21 has the strongest chaperone activity in comparison to RPL7 and ΔC29 Gag, and RPL7 shows better chaperone activity than ΔC29 Gag. DDX21 facilitated tRNA^Pro^ annealing 1.4- and 1.9-fold faster to the WT PBS and 1.7- and 2.2-fold faster to the Δ425-434 PBS compared to RPL7 and ΔC29 Gag, respectively. RPL7 chaperoned tRNA^Pro^ annealing 1.4-fold faster to the WT PBS and 1.3-fold faster to the Δ425-434 PBS compared to ΔC29 Gag (Table 2).

### Both bZIP and CTD of RPL7 are involved in chaperoning the annealing of tRNA^Pro^ to the HTLV-1 PBS

To determine which domains in RPL7 plays a role in tRNA^Pro^ annealing to the PBS, RPL7 variants lacking one or both of the NA-binding domains, ΔbZIP, ΔCTD, or ΔbZIP/CTD, were purified and tested in concentration-dependence annealing assays (Figure S4A and S5). In the presence of 1 μM RPL7 proteins and the less structured Δ425-434 PBS, 46 ± 7% tRNA^Pro^ is annealed by WT, 23 ± 9% by ΔbZIP, 44 ± 5% by ΔCTD, and 12 ± 5% by the ΔbZIP/CTD variant (Figure S5A and S5B). WT and ΔCTD RPL7 have similar chaperone activity, whereas the annealing activity of the ΔbZIP protein is slightly impaired and the activity is further reduced when both bZIP and CTD are deleted. These results indicate that the different RNA binding preferences of bZIP and CTD in RPL7 may together contribute to chaperone tRNA^Pro^ annealing to the PBS and that the bZIP domain plays a more important role in facilitating annealing than the CTD.

### In combinations of two chaperone proteins, only RPL7 and DDX21 synergistically chaperone the annealing of tRNA^Pro^ to the HTLV-1 PBS

To investigate possible synergistic effects of combinations of two proteins, time-course annealing assays were performed in the presence of two chaperone proteins using 0.8 μM DDX21, 0.8 μM RPL7, and 0.8 or 2 μM HTLV-1 WT Gag (Figure S6 – S8). Using the WT PBS domain when only one protein is present, 50 ± 2% of tRNA^Pro^ is annealed by DDX21 (Figure 8G and 8I), 45 ± 3% by RPL7 (Figure 8E and 8I), and 1.52 ± 0.19% by WT Gag after 1 h (Figure 2E and 2I). When 0.8 μM RPL7 and 2 μM HTLV-1 WT Gag are in the same reaction, 37 ± 1% of tRNA^Pro^ is annealed (Figure S6A and S6B). The annealing activity when both RPL7 and Gag proteins are in the reaction is better than WT Gag alone but about the same as RPL7 alone, indicating the absence of a synergistic effect on annealing of tRNA^Pro^ to the WT PBS.

When DDX21 and HTLV-1 WT are in the same reaction, 38 ± 1% of tRNA^Pro^ is annealed by 0.8 μM DDX21 and 0.8 μM WT Gag, and 29 ± 0.2% of tRNA^Pro^ is annealed by 0.8 μM DDX21 and 2 μM WT Gag (Figure S7A – S7C). The annealing activity when both DDX21 and WT Gag proteins are in the reaction is even worse than DDX21 alone, indicating the absence of a synergistic effect on the annealing of tRNA^Pro^ to the WT PBS and the possible competitive binding of these proteins to the substrate.

When 0.8 μM RPL7 and 0.8 μM DDX21 are in the same reaction, 60 ± 2% of tRNA^Pro^ is annealed by RPL7 and DDX21 and the scaled annealing rate (k’) is 0.055 ± 0.008 min^-1^ (Figure S8A and S8B; Table 2). The annealing activity when both RPL7 and DDX21 proteins are in the reaction is 2.1- and 1.5-fold faster than RPL7 or DDX21 alone, respectively, indicating a synergistic effect on the annealing of tRNA^Pro^ to the WT PBS (Table 2).

### When RPL7, DDX21, and HTLV-1 Gag are all present, the annealing of tRNA^Pro^ to the HTLV-1 PBS is more efficient than with a single protein or combinations of two proteins

To investigate possible synergistic effects when all three chaperone proteins are in the same reaction, time-course annealing assays were performed in the presence of 0.8 μM RPL7, 0.8 μM DDX21, and 2 μM HTLV-1 WT Gag (Figure 9A and 9B). Using the WT PBS when three proteins are in the reaction, 54 ± 1% of tRNA^Pro^ is annealed after 1 h and the scaled annealing rate (k’) is 0.086 ± 0.013 min^-1^ (Figure 9A and 9B; Table 2). The annealing activity when three proteins are in the reaction is 1.6-fold faster than the combination of RPL7 and DDX21, and 3.3- and 2.4-fold faster than RPL7 or DDX21 alone, respectively. Thurs, the maximal synergistic effect on the annealing of tRNA^Pro^ to the WT PBS is observed with the combination of RPL7, DDX21, and HTLV-1 Gag (Table 2).

**Figure 9.**
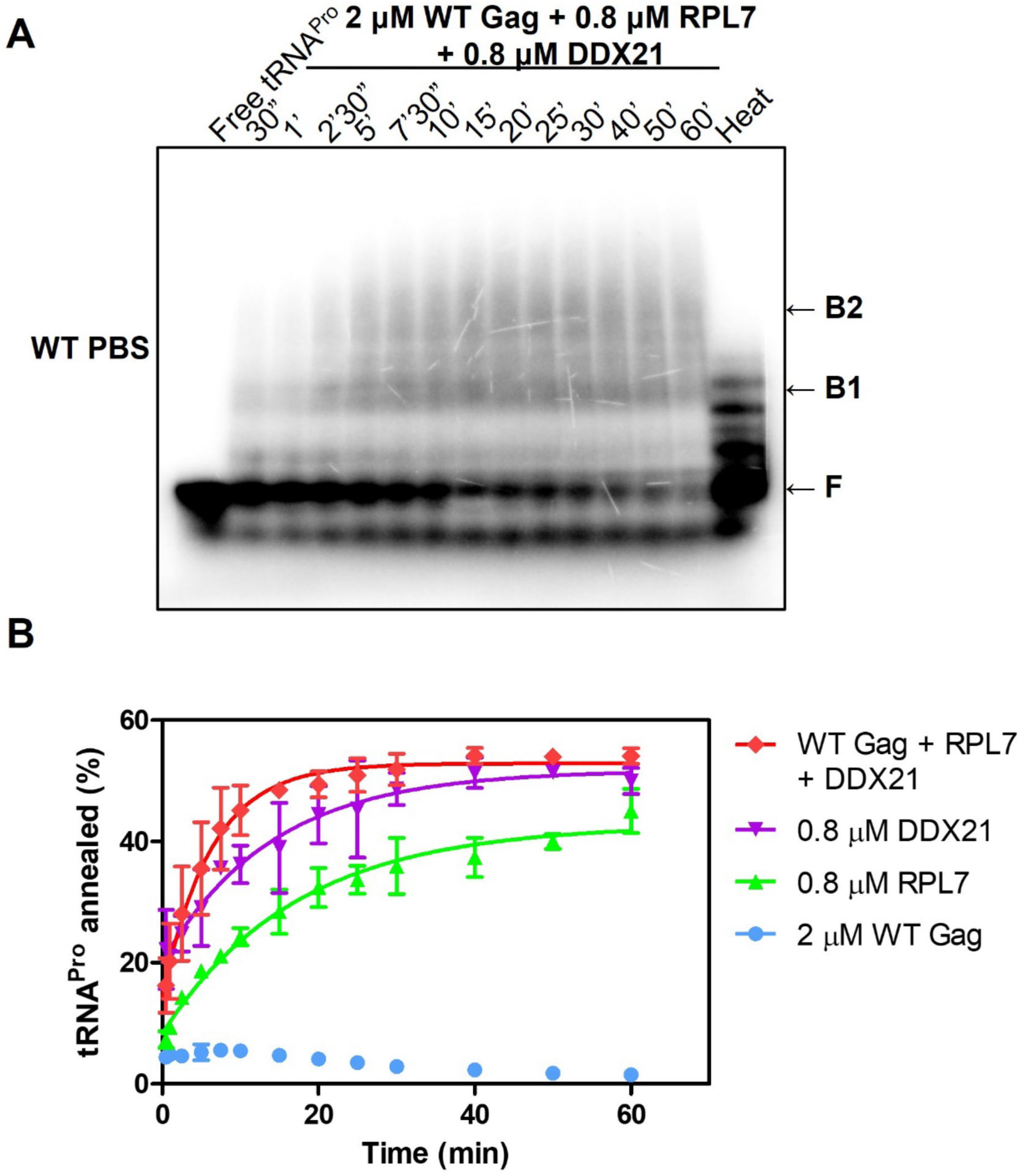
HTLV-1 Gag, RPL7, and DDX21 act synergistically to chaperone tRNA^Pro^ annealing to the HTLV-1 WT PBS. (A) Time-course annealing assays using 20 nM 5’ ^32^P-labeled tRNA^Pro^ and 200 nM HTLV-1 WT PBS in the presence of 2 μM HTLV-1 WT Gag, 0.8 μM RPL7, and 0.8 μM DDX21 at 37°C for varying time. F indicates free tRNA^Pro^ and B1 and B2 indicate different tRNA^Pro^-PBS binary complexes. Heat indicates heat annealing, the positive control. (B) Graph for percentages of tRNA^Pro^ annealed to HTLV-1 WT PBS at different time points in the presence of 2 μM HTLV-1 WT Gag, 0.8 μM RPL7, and 0.8 μM DDX21. Lines represent exponential fits of the data with the standard deviation between trials indicated.

Based on these data, our working model for chaperone-facilitated tRNA^Pro^ annealing to the HTLV-1 PBS is shown in Figure 10. HTLV-1 Gag interacts with RPL7 and DDX21 to form a complex in the host cell and these host factors are packaged into viral particles. tRNA^Pro^ is annealed to the highly-structured HTLV-1 PBS via a two-step mechanism. In the first step, DDX21, a helicase, unwinds the PBS stem-loop. In the second step, RPL7, DDX21, and HTLV-1 Gag synergistically anneal tRNA^Pro^ to the more accessible PBS.

**Figure 10.**
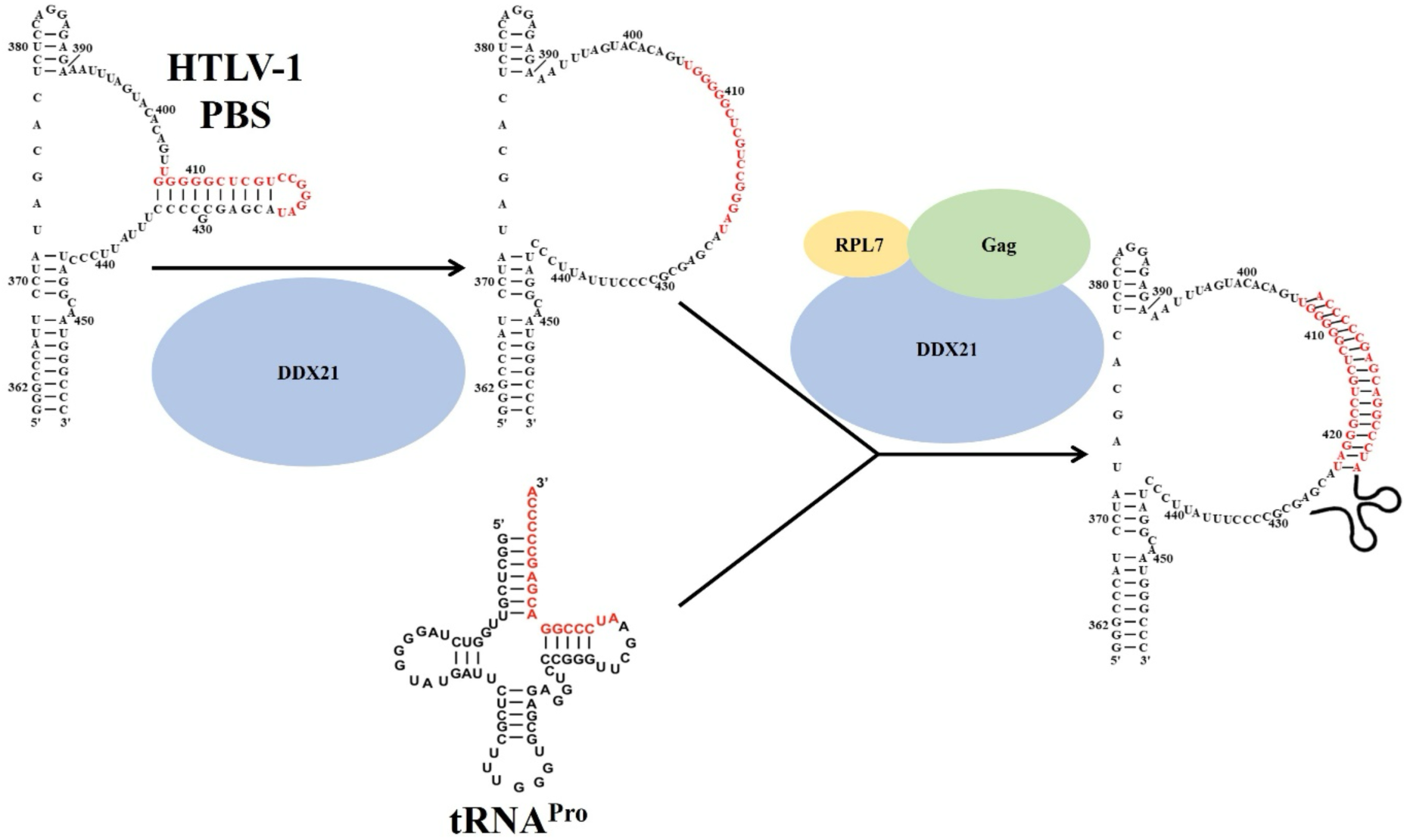
Model for HTLV-1 Gag, human RPL7, and human DDX21 synergistically facilitating tRNA^Pro^ annealing to the HTLV-1 PBS in two steps. In step 1, DDX21 unwinds the stable hairpin in the HTLV-1 PBS region to make the PBS less structured. In step 2, HTLV-1 Gag, RPL7, and DDX21 synergistically facilitate tRNA^Pro^ annealing to the less structured PBS.

## Discussion

Both MA and NC domains of retroviral Gag polyproteins have been reported to possess NA chaperone activity (3,7,16,17,49,50). In HIV-1, the NC domain exhibits stronger chaperone activity than Gag and the presence of MA inhibits Gag’s tRNA^Lys3^ annealing activity (51). While HIV-1 Gag is capable of chaperoning tRNA annealing *in vitro* and this activity depends on the NC domain (51), a two-step annealing was proposed in cells. In this mechanism, Gag facilitates partial tRNA^Lys3^ annealing to the PBS in the host cell, and the mature NC domain promotes formation of more complete tRNA^Lys3^-gRNA interaction in virions (52). HIV-2 Gag has better tRNA^Lys3^ annealing activity *in vitro* than mature MA and NC domains and MA displays stronger annealing activity than NC (50).

In HTLV-1, MA is a more robust NA binding and chaperone protein than NC (16). In this earlier study, the annealing between HIV-1 TAR RNA and DNA was tested but tRNA annealing to the PBS was not investigated. We now show that neither HTLV-1 Gag, NC, or MA can chaperone the annealing of tRNA to the highly-structured PBS domain in which the PBS sequence is embedded in a stable hairpin. Even the robust HIV-1 NC protein failed to significantly facilitate this reaction (data not shown).

In this work, we found that HTLV-1 Gag interacts with RPL7 and DDX21 to form a complex (Figure 3), which increases chaperone activity in a synergistic fashion, with the presence of all three proteins showing more robust activity than any combination of two proteins (Figure 9, S6 - S8). The stoichiometry of the Gag/RPL7/DDX21 complex is unknown. While we showed that HTLV-1 Gag is monomeric in solution (Figure S1B), both RPL7 and DDX21 are capable of forming dimers. RPL7 dimerizes through its N-terminal bZIP domain (42,53), whereas DDX21 forms dimers through the putative hydrophobic dimerization domain (residues 568-620), which is immediately downstream of the helicase core (54). Dimer formation of DDX21 is required for its ATP-dependent double-stranded RNA (dsRNA) unwinding activity and ATP-independent RNA G-quadruplex (RG4) resolving activity (54). Future studies are needed to determine the optimal stoichiometry of the complex for tRNA annealing.

Human RPL7 a ribosomal protein, exhibits tRNA^Pro^ primer annealing activity on its own and both N-terminal bZIP and C-terminal NA-binding domains are involved in this activity (Figure 8 and S5). Human RPL7 is located on the surface of the large 60S subunit of ribosomes and interacts with 28S rRNA (55). In addition to being building blocks of ribosomes and modulating mRNA translation, numerous ribosomal proteins have been reported to have extra-ribosomal functions, such as RNA chaperone activity. In *E. coli*, around a third of 34 tested large ribosomal proteins showed RNA chaperone activity and were proposed to prevent RNAs from being trapped in misfolded forms during translation (56). In HIV-1-infected cells, RPL7 was shown to interact with the NC domain of HIV-1 Gag and act synergistically to increase the chaperone activity of HIV-1 Gag to facilitate annealing of tRNA^Lys3^ to the HIV-1 PBS (21,22). Mouse RPL4 dramatically increases Gag-Pol readthrough in Moloney murine leukemia virus (MoMLV) and HIV-1 possibly through stabilizing the pseudoknot conformation of the frameshift element (57).

Human DDX21, a helicase, also facilitated tRNA^Pro^ annealing to the HTLV-1 PBS on its own and in the absence of ATP (Figure 8). Of the various proteins, tested, DDX21 had the more robust tRNA^Pro^ annealing capability. Helicases that have been reported to be involved in the retroviral lifecycle include RHA (DHX9), DDX1, DDX3, DDX5, DDX17, DDX21, and Moloney leukemia virus 10 homologue (MOV10) (58,59). In HIV-1, DDX21 interacts with both Rev and RRE and facilitates Rev oligomerization on the RRE (35). Other helicases, such as DDX1, DDX3, DDX5, and DDX17, also facilitate the nuclear export of Rev/RRE-dependent viral unspliced and partially spliced transcripts (35,60–63). Although similar functions for these helicases have been reported, knockdowns of individual helicase proteins resulted in distinct negative effects on HIV-1 replication, suggesting that they don’t play redundant roles in modulating the Rev/RRE pathway (62). Whether helicases other than DDX21 and RHA (24) facilitate tRNA RT primer annealing in retroviruses remains to be determined. Whether these helicases are also involved in the HTLV-1 Rex/RexRE-mediated nuclear export of unspliced and partially spliced transcripts is another open question.

Both RPL7 and DDX21 have nuclear localization signals (NLS). RPL7 possesses a tripartite NLS in the bZIP domain and a bipartite NLS in the middle globular domain (64). Similar NLS features are found in most eukaryotic ribosomal proteins, enabling them to be imported into the nucleus for assembly with rRNA into ribosomes in the nucleolus (65). DDX21 is located primarily in the nucleus. In the nucleolus, DDX21 participates in several steps of rRNA biogenesis, including transcription, processing, and small nucleolar ribonucleoprotein (snoRNP)-dependent 2’-O-methylation (66). Upon virus infection, such as vesicular stomatitis virus (VSV) and herpes simplex virus 1 (HSV-1), DDX21 translocates from the nucleus to the cytoplasm to sense viral nucleic acids (67). We showed that both RPL7 and DDX21 are packaged into HTLV-1 virions (Figure 7). Both RPL7 and DDX21 can be present in the nucleus, cytoplasm, and virions, but the location of tRNA primer annealing for any retrovirus remains an open question. The co-packaging of RPL7 and DDX21 along with Gag, gRNA, and tRNA^Pro^ into HTLV-1 virions may ensure that tRNA^Pro^ placement onto the PBS is maintained upon transport from the cytoplasm into virions.

In summary, two HTLV-1 Gag host co-factors, RPL7 and DDX21, were identified in this work and shown to cooperate with Gag to facilitate tRNA^Pro^ annealing to the highly structured HTLV-1 PBS *in vitro.* Investigating how knockdown of RPL7 or DDX21 influences RT primer annealing, other steps of RT, such as elongation and strand transfer, viral production, and HTLV-1 infectivity is ongoing. Targeting the critical primer annealing step by inhibiting RPL7 or DDX21 RNA chaperone activity and Gag-RPL7 or Gag-DDX21 interaction may be a promising therapeutic strategy.

## Materials and Methods

### Plasmid construction

C-terminal 6xHis-tagged human RPL7 bacterial expression plasmid, pPB RPL7 (Figure 1E), was purchased from Applied Biological Materials Inc. N-terminal hemagglutinin (HA)-tagged (YPYDVPDYA) human RPL7 mammalian expression vector, pCMV3 HA-RPL7 (Figure 1F), was purchased from Sino Biological. Bacterial expression plasmid for purifying human DDX21, pMBPDDX21 (Figure 1G), was a gift from Dr. James Williamson at Scripps Research Institute (35). N-terminal mCherry and C-terminal V5-tagged DDX21 mammalian expression vector, pLV CMV mCherry-DDX21-V5 (Figure 1H), was a gift from Dr. Larry Gerace at Scripps Research Institute (Addgene plasmid no.175163) (35).

To prepare a bacterial expression plasmid for purifying HTLV-1 Gag, the coding sequence of HTLV-1 Gag from N3 HTLV-1 Gag (Dr. Louis Mansky, University of Minnesota) was cloned into the backbone of pET3xc HIV-1 Gag (full-length with p6 domain, Dr. Alan Rein, National Cancer Institute). This plasmid, pET3xc HTLV-1 Gag, contains, from the N-terminus, coding regions for HTLV-1 Gag, a Tobacco Etch Virus (TEV) cleavage site (ENLYFQG), and a 6xHis-tag (Figure 1C). To build a mammalian expression construct for HTLV-1 Gag, the coding sequence of HTLV-1 Gag was cut by NheI and BamHI from N3 HTLV-1 Gag and inserted between NheI and BamHI sites in pUCOgs v2.0-1 (Dr. Eddy Arnold, Rutgers University). This construct, pUCOgs HTLV-1 Gag-GFP (green fluorescent protein), contains, from the N-terminus, coding regions for HTLV-1 Gag, human rhinovirus (HRV) 3C protease cleavage site (LEVLFQGP), GFP, and twin-strep tag (Figure 1D). The plasmid for C-terminal FLAG-tagged HTLV-1 Gag (pUCOgs HTLV-1 Gag-FLAG) is derived from pUCOgs HTLV-1 Gag-GFP by swapping HRV 3C site-GFP-twin-strep tag with FLAG tag (DYKDDDDK) using NEB builder HiFi DNA assembly kit (New England Biolabs) according to the manufacturer’s instructions. All constructs for mutants of HTLV-1 Gag, RPL7, and DDX21 were also established by NEB builder HiFi DNA assembly kit (Table S1).

Plasmid templates for *in vitro* transcribing a portion (nt 362-456) of the HTLV-1 5’ UTR including the PBS region (pUC19 HTLV-1 PBS) and human tRNA^Pro^ (pUC119 tRNA^Pro^) were described previously (18,68). To make the HTLV-1 PBS region less structured, nt 425-434 were deleted in pUC19 HTLV-1 PBS using site-directed ligase-independent mutagenesis (SLIM) (Table S1) (69). The mutant construct was named pUC19 HTLV-1 Δ425-434 PBS.

### Protein expression and purification

C-terminal 6xHis-tagged WT and ΔC29 HTLV-1 Gag were expressed in *Escherichia coli* (*E. coli*) Rosetta (DE3). The culture was grown in autoinduction medium (1% tryptone, 0.5% yeast extract, 0.5% glycerol, 0.05% glucose, 0.2% α-lactose, 25 mM Na_2_HPO_4_, 25 mM KH_2_PO_4_, 50 mM NH_4_Cl, 5 mM Na_2_SO_4_, and 2 mM MgSO_4_) from an OD_600_ of 0.05 to 0.6 at 37°C and then the protein of interest was auto-induced at 18°C for 28 h. Cell pellets were lyzed by sonication in lysis buffer (20 mM Tris-HCl, pH 7.4, 500 mM NaCl, 1 μM ZnCl_2_, 5 mM 2-mercaptoethanol (β-ME), 10% glycerol, 0.05% Triton X-100, and tablets of protease inhibitor (Roche)). Polyethylenimine (PEI) was added to the soluble fraction to a final concentration of 0.6% (v/v) to precipitate and remove nucleic acids, and ammonium sulfate was then added to the supernatant to a final concentration of 1.3 M. Ammonium sulfate-containing pellets were resuspended in the buffer A (20 mM Tris-HCl, pH 7.4, 500 mM NaCl, 1 μM ZnCl_2_, and 5 mM β-ME), and then applied to a HIS-Select nickel affinity column (Sigma-Aldrich). The column was washed with buffer A supplemented with 5 mM imidazole and eluted with a step gradient of 10 mM, 20 mM, 50 mM, 75 mM, 100 mM, 150 mM, and 200 mM imidazole. All fractions were analyzed by sodium dodecyl sulfate-polyacrylamide gel electrophoresis (SDS-PAGE) followed by Coomassie blue staining. The elution fractions containing the most concentrated protein of interest were pooled and dialyzed into buffer A to remove imidazole at 4°C overnight. The 6xHis tag was cleaved using 3 mg of TEV protease per 20 mg of HTLV-1 Gag during the dialysis. After overnight dialysis and TEV cleavage, the sample was loaded onto a HIS-Select nickel affinity column (Sigma-Aldrich) again to remove the cleaved 6xHis tag and 6xHis-tagged

TEV protease. The NaCl concentration of the fractions containing the purest protein of interest was reduced from 500 mM to 100 mM by 5-fold dilution with the buffer B (20 mM Tris-HCl, pH 7.4, 1 μM ZnCl_2_, and 5 mM β-ME) and applied to a 5 mL HiTrap heparin column (Cytiva). The column was washed with 3 column volumes (CV) of buffer B with 0.5 M NaCl and eluted with 4 CV of buffer B with 1 M NaCl. The fractions containing the purest HTLV-1 Gag were combined and dialyzed into buffer A supplemented with 10% glycerol. The protein concentration of HTLV-1 Gag was determined by measuring the absorbance at 280 nm and using a molar extinction coefficient of 60390 M^-1^cm^-1^.

C-terminal 6xHis-tagged human RPL7 was expressed in *E. coli* Rosetta (DE3). The culture was grown in Luria-Bertani (LB) broth (1% tryptone, 0.5% yeast extract, and 0.5% NaCl) from an OD_600_ of 0.05 to 0.6 at 37°C and the protein of interest was induced with 1 mM of isopropyl β-D-1-thiogalactopyranoside (IPTG) at 37°C for 24 h. The protein was purified as previously described with some modifications (70). Briefly, cell pellets were dissolved in the denaturing lysis buffer (25 mM Tris-HCl, pH 8.0, 100 mM NaCl, 6 M guanidine hydrochloride, 0.5 mM phenylmethylsulfonyl fluoride (PMSF), and tablets of protease inhibitor (Roche)) to release RPL7 protein from inclusion bodies. The soluble fraction was loaded onto a HIS-Select nickel column (Sigma-Aldrich). The column was washed with 8 CV of denaturing lysis buffer and 8 CV of urea buffer (25 mM Tris-HCl, pH 8.0, 100 mM NaCl, 8 M urea, 10 mM imidazole, and 0.5 mM PMSF), and eluted with 15 CV of urea buffer supplemented with 100 mM of imidazole. The elution fractions containing the purest RPL7 were pooled and dialyzed into 25 mM Tris-HCl, pH 8.0, 150 mM NaCl, 5 mM β-ME, and 10% glycerol. The protein concentrations of RPL7 WT and truncation mutants were examined by measuring the absorbance at 280 nm and using the following molar extinction coefficients: WT, 33350 M^-1^cm^-1^; ΔbZIP (ΔN77), 27390 M^-1^cm^-1^; ΔCTD (ΔC51) 27850 M^-1^cm^-1^; ΔbZIP/CTD, 21890 M^-1^cm^-1^.

N-terminal maltose-binding protein (MBP) and C-terminal 6xHis-tagged human DDX21 was expressed in *E. coli* Rosetta (DE3). The culture was grown in LB broth from an OD_600_ of 0.05 to 0.6 at 37°C and then the protein of interest was induced with 1 mM of IPTG at 18°C for 20 h. The protein was purified as previously established protocol with some alterations (35).

Briefly, cells were lyzed by sonication in lysis buffer (20 mM Tris-HCl, pH 7.8, 500 mM NaCl, 1 mM ethylenediaminetetraacetic acid (EDTA), and 1 mM Tris(2-carboxyethyl)phosphine (TCEP), 10% glycerol, 0.05% Triton X-100, and tablets of protease inhibitor (Roche)). PEI was added to the soluble fraction to a final concentration of 0.025% (v/v) to precipitate and remove nucleic acids, and the clear supernatant was applied to a HIS-Select nickel affinity column (Sigma-Aldrich). The column was washed with buffer A (20 mM Tris-HCl, pH 7.8, 500 mM NaCl, 1 mM EDTA, 1 mM TCEP, and 2% glycerol) supplemented with 5 mM imidazole and eluted with a step gradient of 10 mM, 20 mM, 50 mM, 75 mM, 100 mM, 150 mM, and 200 mM imidazole. The elution fractions containing the most concentrated protein of interest were pooled and dialyzed into buffer A to remove imidazole at 4°C overnight. The MBP tag was cleaved using 3 mg of TEV protease per 20 mg of DDX21 during dialysis. After overnight dialysis and TEV cleavage, the NaCl concentration of the sample was reduced from 500 mM to 75 mM by 6.7-fold dilution with the buffer B (20 mM Tris-HCl, pH 7.8, 1 mM EDTA, 1 mM TCEP, and 2% glycerol) and loaded onto a 5 mL HiTrap heparin column (Cytiva). The column was washed with 3 CV of buffer B with 0.5 M NaCl and eluted with 4 CV of buffer B with 1 M NaCl and 3 CV of buffer B with 2 M NaCl. The fractions containing the purest DDX21 were combined and dialyzed into 20 mM Tris-HCl, pH 7.8, 500 mM NaCl, 1 mM TCEP, and 10% glycerol. The protein concentration of DDX21 was determined from the absorbance at 280 nm and using a molar extinction coefficient of 48360 M^-1^cm^-1^.

### RNA preparation

Plasmids encoding HTLV-1 PBS region and human tRNA^’()^ driven by a T7 promoter with a PstI or FokI site at the 3’ end were linearized and used as a template for *in vitro* transcription using T7 RNA polymerase as previously described (Figure 1A and 1B; Table S2) (71). *In vitro* transcribed RNA was loaded onto a 10% polyacrylamide/8 M urea gel, and the gel piece containing the RNA of interest was excised and eluted in RNA elution buffer (0.5 mM NH_4_OAc and 1 mM EDTA, pH 8.0) at 37°C overnight. The gel eluate was concentrated by butanol extraction and ethanol precipitation. The concentration of RNAs was calculated according to Beer’s law by measuring the absorbance at 260 nm and using the following molar extinction coefficients: HTLV-1 WT PBS, 870278 M^-1^cm^-1^; HTLV-1 Δ425-434 PBS, 776828 M^-1^cm^-1^; tRNA^’()^, 655343 M^-1^cm^-1^.

Prior to use, all RNAs were folded in 50 mM 4-(2-hydroxyethyl)-1-piperazineethanesulfonic acid (HEPES), pH 7.5 at 80°C for 2 min, 60°C for 2 min, addition of MgCl_2_ to 1 mM, incubation at 37°C for 30 min, and cooling on ice for at least 30 min.

### 5’ ^32^P RNA labeling

To remove the 5’ phosphate before labeling, 500 pmol of *in vitro* transcribed RNAs was incubated with 5 units of calf intestinal alkaline phosphatase (CIP) (New England Biolabs) at 37°C for 1 h. After phenol-chloroform extraction and ethanol precipitation, RNAs were treated with 50 μCi of [γ-^32^P] ATP (PerkinElmer) and 10 units of T4 polynucleotide kinase (PNK) (New England Biolabs) to phosphorylate the 5’ end. Free [γ-^32^P] ATP was cleaned up by a sephadex G-25 spin column (Roche) and labeled RNAs were recovered by phenol-chloroform extraction and ethanol precipitation.

### Gel-shift tRNA annealing assays

HTLV-1 WT (98 nt) or Δ425-434 PBS (88 nt) and 5’ ^32^P-labeled tRNA^’()^ (75 nt) were folded separately as described in RNA preparation. Folded HTLV-1 PBS and tRNA^’()^ were then incubated in the same reaction at 37°C for 10 min and cooled to room temperature before the addition of chaperone proteins (Figure 1A). After adding chaperone proteins, such as HTLV-1 Gag, human RPL7, and/or human DDX21, the final reaction mixture (10 μL) contained 20 nM 5’ ^32^P-labeled tRNA^’()^, 200 nM HTLV-1 PBS, 50 mM HEPES, pH 7.5, 150 mM NaCl, 1 mM MgCl_2_, 0.1 mM ZnCl_2_, 5 mM dithiothreitol (DTT), and 0-4 μM chaperone proteins. In single-time-point concentration-dependence assays, the reaction mixtures containing different protein concentrations were incubated at 37°C for 1 h. In time-course kinetic assays, the reaction aliquots with a single protein concentration, 0.8 μM RPL7, 0.8 μM DDX21, and/or 2 μM HTLV-1 Gag, were taken at the indicated time point until 1 h at 37°C. For heat annealing (the positive control), HTLV-1 PBS and tRNA^’()^ were folded together as described in RNA preparation. In all annealing assays, reactions were quenched with 1% SDS and 1 mg/ml proteinase K (New England Biolabs) and incubated at 37°C for 30 min. After phenol-chloroform extraction, the samples were mixed with 6x native loading dye (50% glycerol, 0.25% bromophenol blue, 0.25% xylene cyanol) and loaded onto pre-cast NativePAGE Bis-Tris 4-16% polyacrylamide gradient gels (Invitrogen). The running buffer was 1x TB (89 mM Tris-HCl and 89 mM boric acid, pH 8.3) supplemented with 1 mM MgCl_2_. Gels were exposed to phosphor screens overnight and bands were visualized with Typhoon FLA 9500 (Cytiva) and quantified using Image J software. Time-course annealing data were fit to the following single-exponential equation using GraphPad Prism software: Annealed(t) = Annealed_max_ – (Annealed_max_-Annealed_min_) X e^-kt^, where Annealed (t) is the fraction tRNA^’()^ annealed as a function of time and k is the annealing rate. Concentration-dependence annealing data were fit to the following equation: Annealed (C) = (Annealed_max_ X C) / (K_1/2_ + C), where Annealed (C) is the fraction tRNA^’()^ annealed as a function of protein concentration and K_1/2_ is the protein concentration when the annealing is half maximal.

### Cell culture and virus fractionation

MT-2 cells were cultured in Roswell Park Memorial Institute (RPMI) 1640 medium supplemented with 10% (vol/vol) fetal bovine serum (FBS), 100 unit/mL penicillin, and 100 μg/mL streptomycin. Human embryonic kidney cells (HEK293T) were grown in Dulbecco’s Modified Eagle Medium (DMEM) supplemented with 10% FBS, 100 unit/mL penicillin, 100 μg/mL streptomycin, and 1x non-essential amino acids solution (complete DMEM). All cells are maintained in a 37°C incubator with 5% CO_2_.

To collect HTLV-1 virions, 10 mL of culture supernatant from MT-2 cells was filtered through 0.45-μm syringe filters, layered on the top of 1 mL of 25% sucrose, and concentrated by ultracentrifugation at 90,000 x g for 1.5 h at 4°C in a Sorvall SW41 swinging bucket rotor (sucrose cushion). The viral pellet was resuspended in 500 μL of 1x STE buffer (10 mM Tris-HCl, pH 8.0, 100 mM NaCl, and 1 mM EDTA) by gentle shaking at 4°C overnight. Resuspended viruses were applied to 10 mL of OptiPrep density gradient medium (iodixanol, Sigma-Aldrich) as 5 steps with 7.5% increments ranging from 10 to 40% and centrifuged at 250,000 x g for 3 h at 4°C in a SW41 rotor. After ultracentrifugation, the sample was divided equally from the top to the bottom into 9 fractions. These 9 fractions were concentrated using sucrose cushion as described above and analyzed by immunoblotting.

### Affinity tagging/purification-mass spectrometry (AP-MS)

One 100-mm dish seeded with 4 million HEK293T cells were transfected with 10 μg of plasmid expressing GFP or HTLV-1 Gag-GFP (Figure 1D) by PEI method next day after seeding and harvested 48 h post-transfection (72). Thirty million cells harvested from three 100-mm dishes were lysed in 1 mL of lysis buffer (100 mM HEPES, pH 8.0, 150 mM NaCl, 0.5% 3-((3-cholamidopropyl) dimethylammonio)-1-propanesulfonate (CHAPS), 1 mM TCEP, 25 mM NaF, and tablets of protease inhibitor (Roche)) with gentle rocking at 4°C for 30 min. Crude cell lysate was cleared of debris via centrifugation at 15,000 x g at 4°C for 30 min. MegStrep XT type3 beads (5 μL, IBA Lifesciences) were added to the clear cell lysate and samples were rotated at 4°C overnight. Beads were washed with 1 mL of 100 mM HEPES, pH 8.0, 150 mM NaCl, and 1 mM TCEP three times, and resuspended in 20 μL of phosphate-buffered saline (PBS, 2.67 mM KCl, 137.93 mM NaCl, 1.47 mM KH_2_PO_4_, 8.06 mM Na_2_HPO_4_, pH 7, Gibco).

One-fourth of the sample was analyzed by SDS-PAGE followed by Coomassie blue staining to check whether our protein of interest was pulled down (Figure S2A). The rest of the sample was further processed by in-bead trypsin proteolysis and analyzed by liquid chromatography with tandem mass spectrometry (LC-MS/MS) peptide sequencing at the Mass Spectrometry and Proteomics (MSP) facility, Campus Chemical Instrument Center (CCIC), Ohio State University.

### Immunoprecipitation (IP) and immunoblotting

HEK293T cells transfected with indicated plasmids by PEI method were harvested 48 h post-transfection. Ten million cells were lysed in 0.5 mL of lysis buffer (PBS supplemented with 1% Triton X-100 and tablets of protease inhibitor (Roche)) with gentle rotating at 4°C for 30 min. Crude cell lysate was centrifuged at 15,000 x g at 4°C for 30 min to remove cell debris. The concentration of protein in the clear cell lysate was determined by Pierce bicinchoninic acid (BCA) protein assay kit (Thermo Scientific). For HTLV-1 Gag-FLAG co-IP, 10 μL of anti-FLAG M2 magnetic beads (Sigma-Aldrich) were added to the cell lysate and samples were rotated at 4°C overnight. Beads were washed with 1 mL of lysis buffer 6 times. For RPL7-HA and DDX21 co-IP, 2 μg of anti-HA (BioLegend), anti-DDX21 (Proteintech), mouse IgG isotype control (Invitrogen, as an HA co-IP background control), or rabbit IgG isotype control (Invitrogen, as a DDX21 co-IP background control) antibody was added to the clear cell lysate and samples were rotated at 4°C overnight. The next day, 25 μL of Dynabeads Protein G (Invitrogen) for RPL7-HA co-IP or protein A (Invitrogen) for DDX21 co-IP were added and the sample was rocked for additional 4 h at 4°C. Beads were washed with 1 mL of lysis buffer 6 times.

For p24 co-IP in MT-2 cells, five million cells were lysed in 0.5 mL of PBS supplemented with 1% Triton and tablets of protease inhibitor (Roche). The clear lysate was incubated and rocked with 2 μg of anti-HTLV-1 p24 antibody (Santa Cruz) or mouse IgG isotype control (Invitrogen) at 4°C overnight. The following day, 25 μL of Dynabeads Protein G (Invitrogen) were added and the sample was rotated at 4°C for additional 4 h. Beads were washed with 0.5 mL PBS three times.

After the washing step, beads were resuspended in 20 μL of lysis buffer with 5 μL of 5x protein loading dye (300 mM Tri-HCl, pH 6.8, 25% Glycerol, 10% SDS, 715 mM β-ME, 0.1% bromophenol blue) and boiled for 10 min before loading onto 10% polyacrylamide denaturing gels or pre-cast NuPAGE Bis-Tris 12% polyacrylamide gels (Invitrogen). Proteins in the gels were transferred onto polyvinylidene fluoride (PVDF) membranes (Cytiva). Membranes were blocked with 5% non-fat milk (Bio-Rad) in 19 mM Tris-HCl, 137 mM NaCl, 27 mM KCl, and 0.1% Tween-20 (TBST) and incubated with primary antibodies at 4°C overnight. The following primary antibodies were used: anti-GFP (Invitrogen, 1:5000), anti-FLAG (Sigma-Aldrich, 1:1000), anti-HA (BioLegend, 1:2000), anti-V5 (Invitrogen, 1:1000), anti-HTLV-1 p19 (ZeptoMetrix, 1:1000), anti-HTLV-1 p24 (Santa Cruz, 1:1000), anti-RPL7 (Bethyl, 1:2000), anti-DDX21 (Proteintech, 1:1000), and anti-β-actin (Sigma-Aldrich, 1:5000). After washed with TBST three times, membranes were incubated with secondary antibodies at room temperature for 1 h. The secondary antibodies used were horseradish peroxidase (HRP)-conjugated goat anti-mouse (Promega, 1:5000) or goat anti-rabbit antibodies (Promega, 1:5000). The blots were developed using SuperSignal West Pico PLUS Chemiluminescent Substrate (Thermo Scientific) or Pierce Enhanced Chemiluminescence (ECL) Western Blotting Substrate (Thermo Scientific) and visualized by Amersham Imager 680 imaging system (Cytiva).

## Acknowledgements

We thank Dr. Sophie Harvey for performing tandem mass spectrometry (LC-MS/MS) peptide sequencing at the Mass Spectrometry and Proteomics (MSP) facility, Campus Chemical Instrument Center (CCIC), Ohio State University. We thank Drs. Patrick L. Green and Amanda R. Panfil (Ohio State University) for providing MT-2 cells and sharing materials, facilities, and lab space to perform cell culture work. We thank Dr. Weixin Wu and Dr. Joshua Hatterschide for generating the pUC19 HTLV-1 WT and Δ425-434 PBS constructs. This work was supported by the National Institute of Allergy and Infectious Disease of the National Institutes of Health (NIH) under award numbers R01AI150493 and U54AI170855. The content is solely the responsibility of the authors and does not necessarily represent the official views of the NIH.

