## Supplementary Files for "Human RPL7 and DDX21 interact with HTLV-1 Gag and enhance tRNA^Pro^ primer annealing to genomic RNA"

### Materials and Methods

#### Circular dichroism (CD) spectroscopy

CD spectra were obtained on a Jasco J-815 spectrometer using 2.5 μM WT RPL7 at 25°C in 50 mM sodium phosphate, pH 7.8, at Campus Chemical Instrument Center (CCIC), Ohio State University. CD absorbance values were recorded from 185 to 275 nm with a data pitch of 0.5 nm at a scanning speed of 100 nm/min. Final CD spectra were plotted against wavelength in nm after subtracting the CD values for the buffer blank.

#### Size-exclusion chromatography with multi-angle laser-light scattering (SEC-MALS)

HTLV-1 WT or ΔC29 Gag proteins (100 μL of 15 μM) were loaded onto a Superdex 200 Increase 10/300 GL size-exclusion column (Cytiva) and eluted with 20 mM Tris-HCl, pH 7.4, 500 mM NaCl, 1 μM $ZnCl_2$, and 5 mM β-mercaptoethanol at a flow rate of 0.5 mL/min. The SEC was controlled by an ÄKTA Pure 25 chromatography system (Cytiva) and MALS analysis was performed by a DAWN HELIOS $8^+$ laser detector (Wyatt technology) and an Optilab T-rEX refractometer (Wyatt technology) conjugated to the SEC control system. The light-scattering (LS) intensity and differential refractive index (dRI) of each eluate fraction were measured by the laser detector and the refractometer, respectively. UV, LS, and dRI signals were aligned and the molecular weight of each signal peak was calculated using ASTRA 7.1.4.8 software.

### Figures and Tables

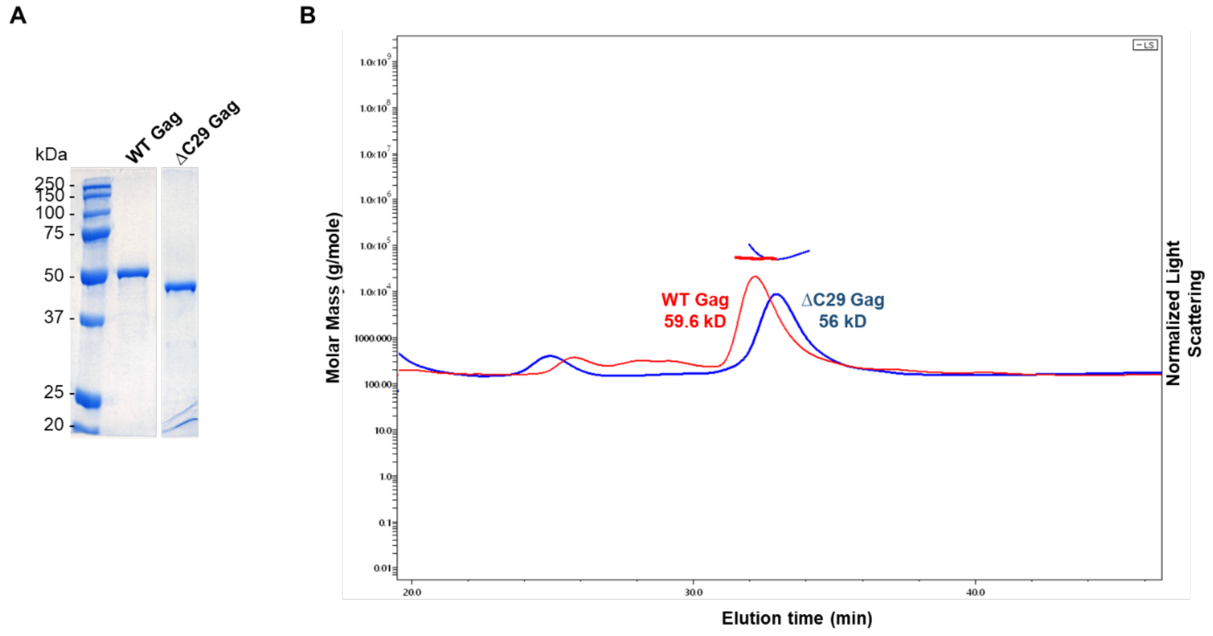


**Figure S1. HTLV-1 Gag recombinant proteins were successfully purified from *E. coli* for the first time and they form monomers in solution inferred from size-exclusion chromatography with multi-angle laser-light scattering (SEC-MALS) analysis.** (A) Recombinant HTLV-1 WT and ΔC29 Gag proteins (1 μg) purified from *E. coli.* were run on an SDS-polyacrylamide (10%) gel and stained with Coomassie blue and only one major band for each protein migrated close to their molecular weights. (B) SEC-MALS chromatogram of purified HTLV-1 Gag proteins. Normalized light scattering peaks for HTLV-1 WT (red) and ΔC29 (blue) Gag are shown. The molar mass range for each peak was calculated and indicated by thick lines.

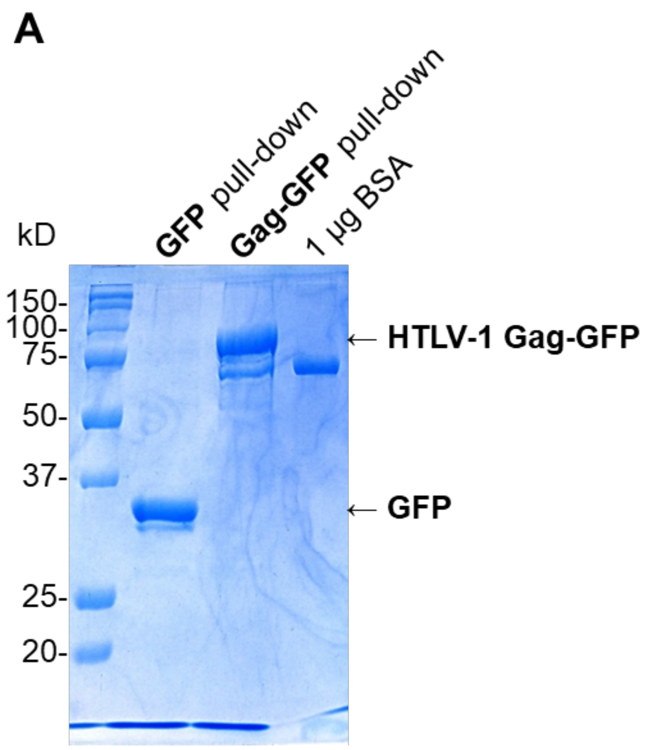


**Figure S2.** Affinity tagging/purification-mass spectrometry (AP-MS) was used to identify interacting partners of HTLV-1 Gag. (A) Prior to AP-MS analysis, GFP or HTLV-1 Gag-GFP pulled-down by streptactin from HEK293T cell lysate 48 h post-transfection was run on an SDS-polyacrylamide gel (10%) and stained with Coomassie blue.

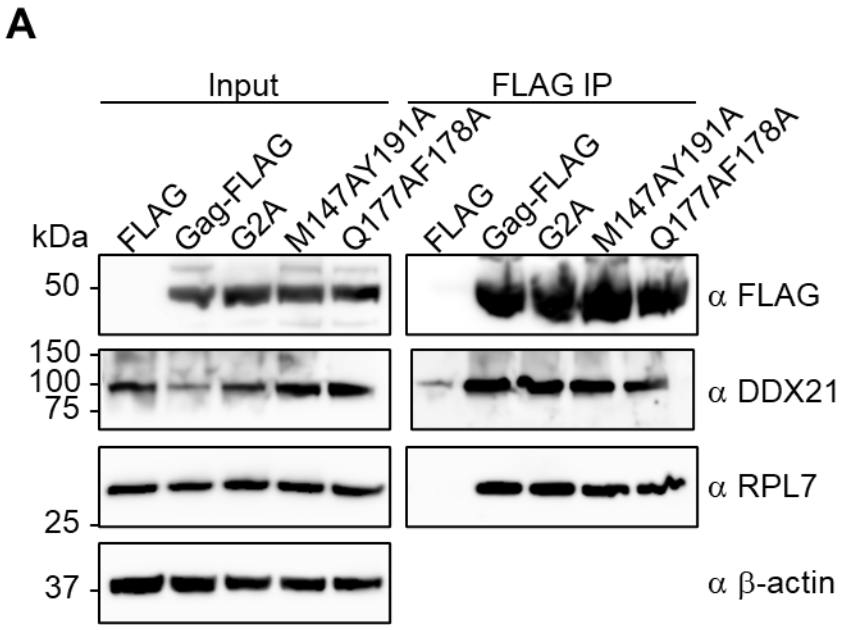


**Figure S3. The interactions of HTLV-1 Gag with RPL7 and DDX21 are Gag-myristoylation- and Gag-oligomerization-independent.** (A) FLAG, and WT, G2A, M147AY191A, and Q177AF178A Gag-FLAG in HEK293T cell lysate were immunoprecipitated and immunoblotted using an anti-FLAG antibody 48 h post-transfection. Co-immunoprecipitated proteins were immunoblotted by anti-DDX21 and anti-RPL7 antibodies. β-actin was used as a loading control.

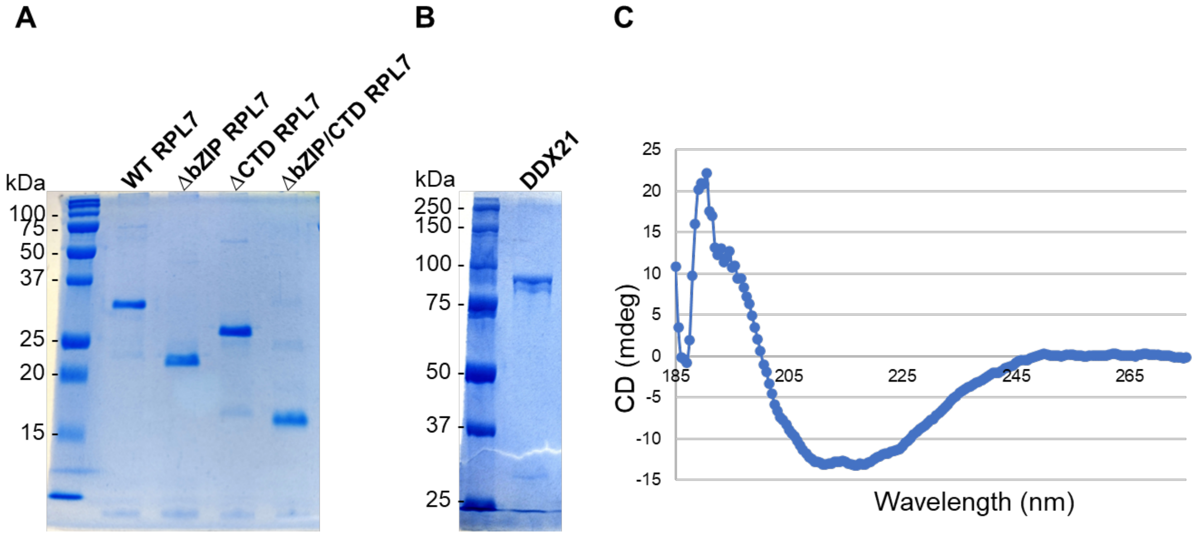


**Figure S4. Analysis of the purity of human WT, ∆basic leucine zipper (bZIP), ∆C-terminal domain (CTD), and ∆bZIP/CTD RPL7, and human WT DDX21 recombinant proteins purified from *E. coli*. by SDS-PAGE.** (A) Recombinant human WT, ∆bZIP, ∆CTD, and ∆bZIP/CTD RPL7 proteins (1 µg) purified from *E. coli.* were run on an SDS-polyacrylamide (12%) gel and stained with Coomassie blue. (B) Recombinant human DDX21 protein (1 µg) purified from *E. coli.* was run on an SDS-polyacrylamide (8%) gel and stained with Coomassie blue. (C) Circular dichroism (CD) spectrum of WT RPL7 showed that recombinant RPL7 protein was well-folded.

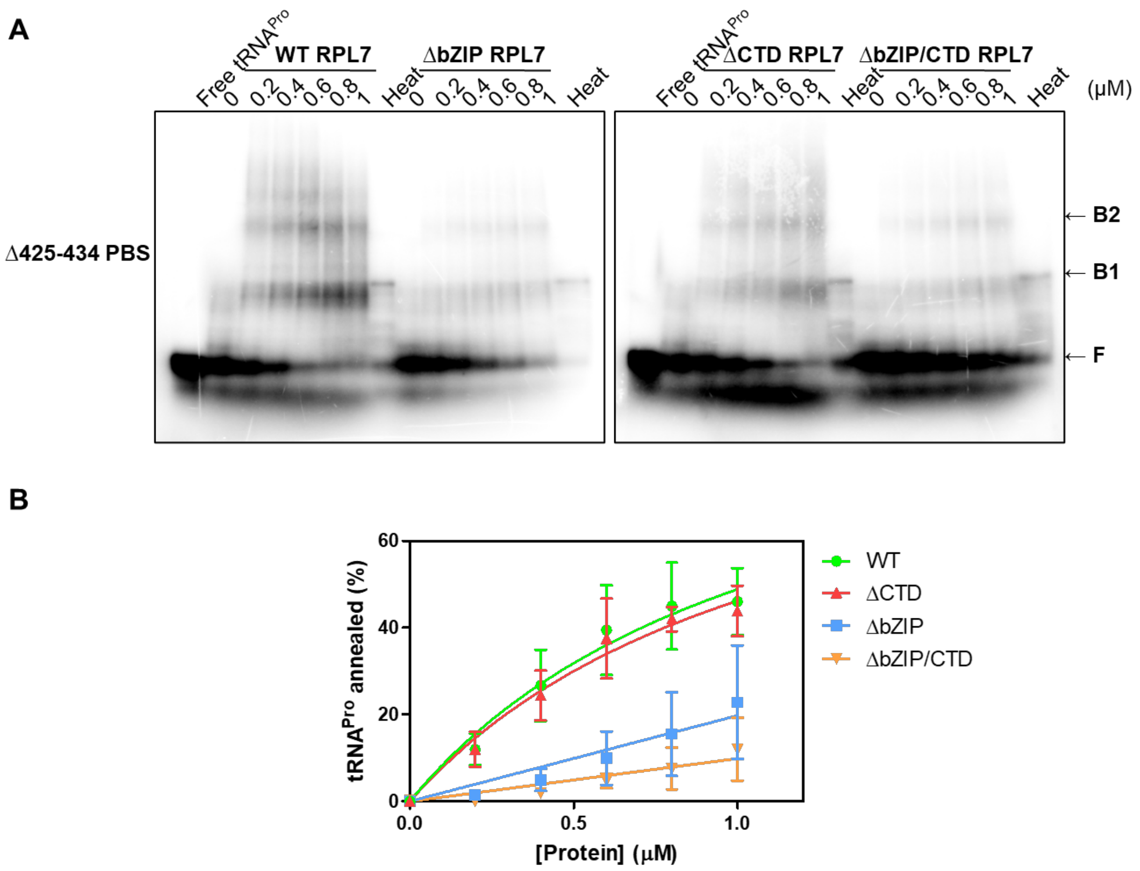


**Figure S5. Both bZIP and CTD of RPL7 are involved in chaperoning tRNA$^{Pro}$ annealing to the HTLV-1 PBS.** (A) Concentration-dependence annealing assays using 20 nM 5' $^{32}$P-labeled tRNA$^{Pro}$ and 200 nM HTLV-1 Δ425-434 PBS in the presence of varying concentrations of WT, ΔbZIP, ΔCTD, or ΔbZIP/CTD RPL7 at 37°C for 1 h. F indicates free tRNA$^{Pro}$, and B1 and B2 indicate different conformations of tRNA$^{Pro}$-PBS binary complexes. Heat indicates heat annealing, the positive control. (B) Graph for percentages of tRNA$^{Pro}$ annealed to the Δ425-434 PBS in varying concentrations of WT, ΔbZIP, ΔCTD, or ΔbZIP/CTD RPL7.

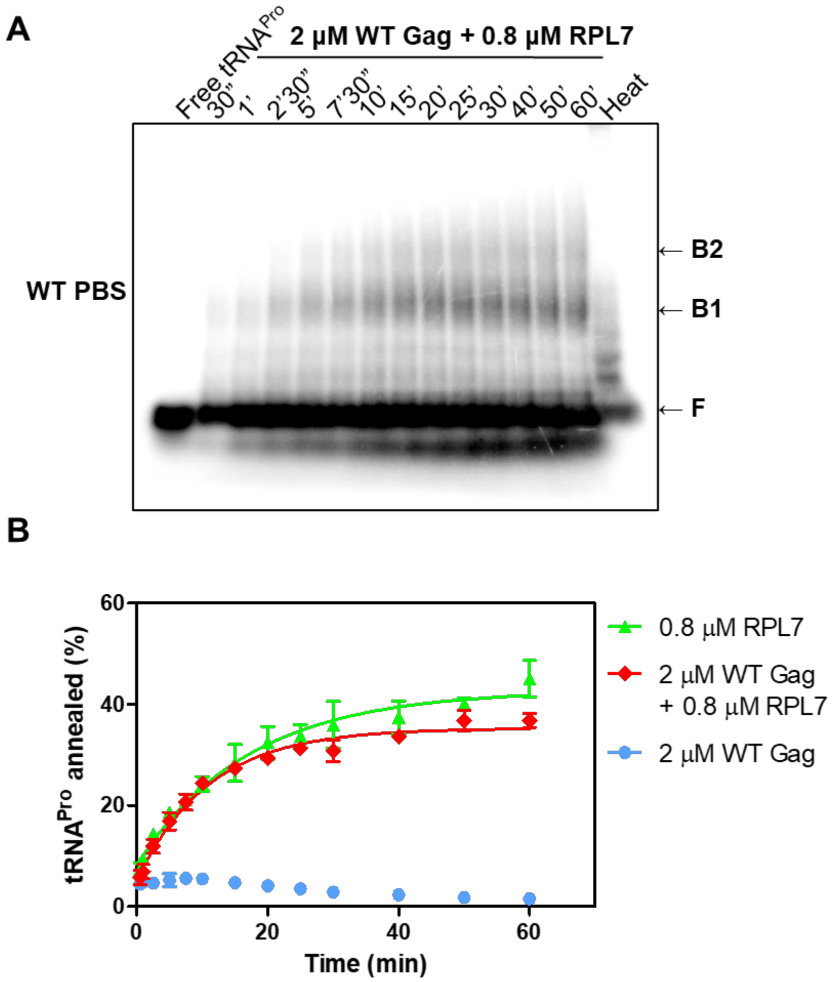


**Figure S6. In the presence of both RPL7 and HTLV-1 Gag, there is no synergistic activity to chaperone tRNA$^{Pro}$ annealing to the HTLV-1 PBS.** (A) Time-course annealing assays using 20 nM 5' $^{32}$P-labeled tRNA$^{Pro}$ and 200 nM HTLV-1 WT PBS in the presence of 0.8 μM RPL7 and 2 μM HTLV-1 WT Gag at 37°C for varying time. F indicates free tRNA$^{Pro}$, and B1 and B2 indicate different tRNA$^{Pro}$-PBS binary complexes. Heat indicates heat annealing, the positive control. (B) Graph showing percentages of tRNA$^{Pro}$ annealed to HTLV-1 WT PBS at different time points in the presence of 0.8 μM RPL7 and 2 μM HTLV-1 WT Gag. Lines represent exponential fits of the data with the standard deviation between trials indicated.

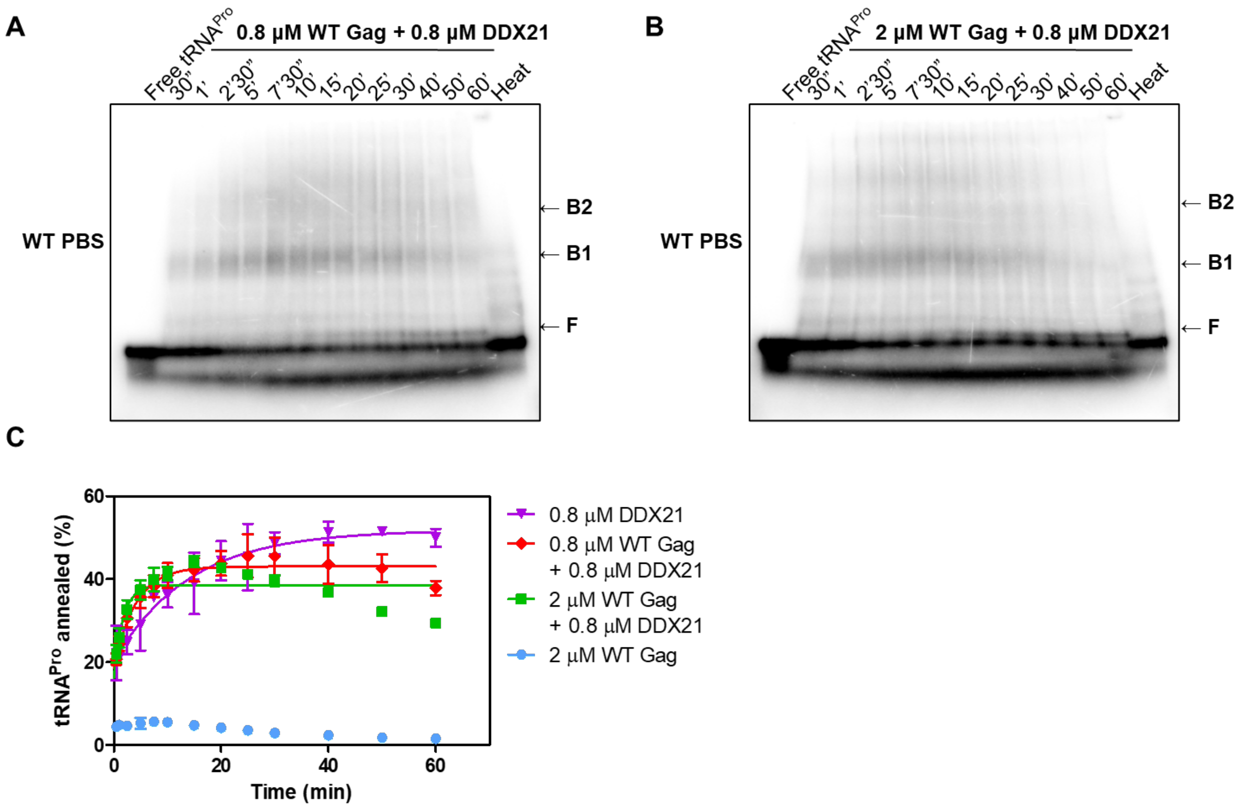


**Figure S7. In the presence of both DDX21 and HTLV-1 WT Gag, there is no synergistic effect to chaperone tRNA$^{Pro}$ annealing to the HTLV-1 PBS.** (A and B) Time-course annealing assays using 20 nM 5' $^{32}$P-labeled tRNA$^{Pro}$ and 200 nM HTLV-1 WT PBS in the presence of 0.8 μM DDX21 and 0.8 μM (A) or 2 μM (B) HTLV-1 WT Gag at 37°C for varying time. F indicates free tRNA$^{Pro}$, and B1 and B2 indicate different conformations of tRNA$^{Pro}$-PBS binary complexes. Heat indicates heat annealing, the positive control. (C) Graph for percentages of tRNA$^{Pro}$ annealed to HTLV-1 WT PBS at different time points in the presence of 0.8 μM DDX21 and 0.8 μM or 2 μM HTLV-1 WT Gag. Lines represent exponential fits of the data with the standard deviation between trials indicated.

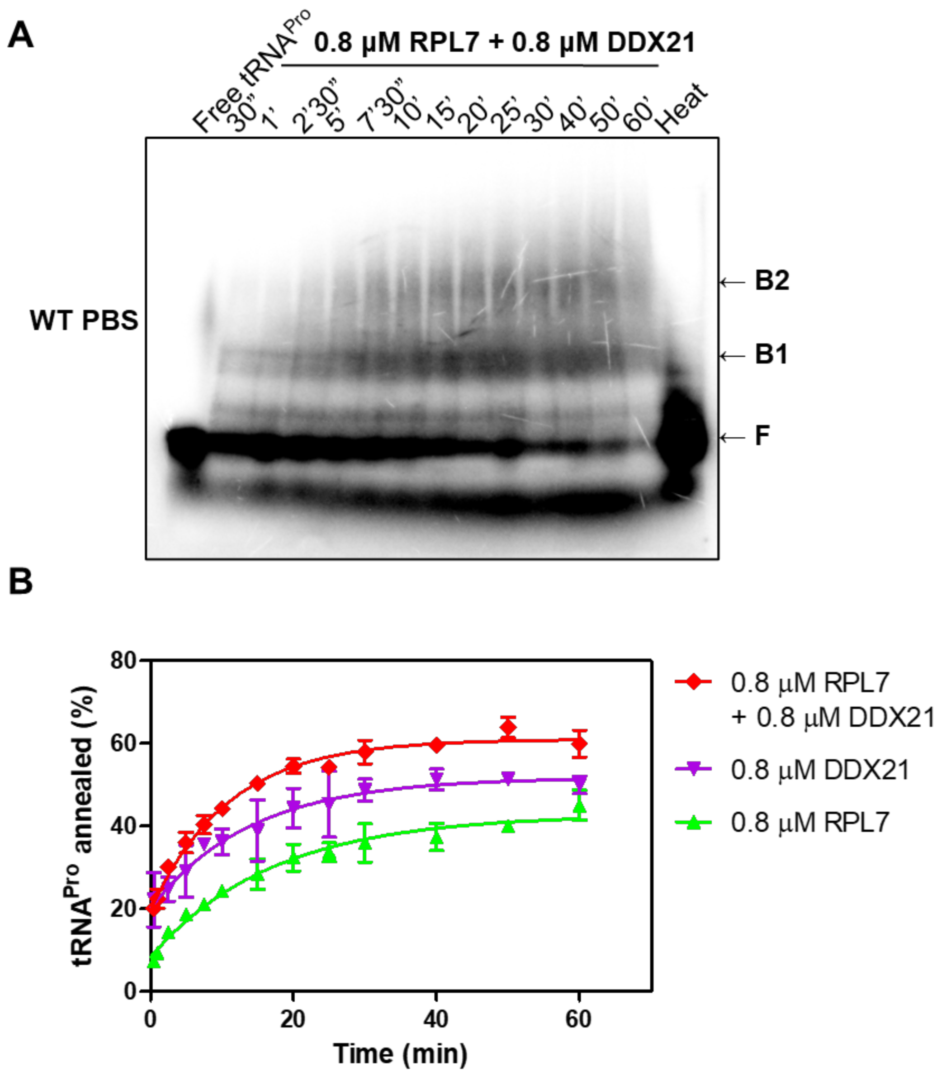


**Figure S8. RPL7 and DDX21 act synergistically to chaperone tRNA$^{Pro}$ annealing to the HTLV-1 PBS.** (A) Time-course annealing assays using 20 nM 5' $^{32}$P-labeled tRNA$^{Pro}$ and 200 nM HTLV-1 WT PBS in the presence of 0.8 μM RPL7 and 0.8 μM DDX21 at 37°C for varying time. F indicates free tRNA$^{Pro}$, and B1 and B2 indicate different tRNA$^{Pro}$-PBS binary complexes. Heat indicates heat annealing, the positive control. (B) Graph for percentages of tRNA$^{Pro}$ annealed to HTLV-1 WT PBS at different time points in the presence of 0.8 μM RPL7 and 0.8 μM DDX21. Lines represent exponential fits of the data with the standard deviation between trials indicated.

**Table S1.** List of primer sets used for generating pET3xc HTLV-1 Gag, pUCOgs HTLV-1 Gag-GFP, pUCOgs HTLV-1 Gag-FLAG, pPB RPL7, pCMV3 HA-RPL7, pLV CMV mCherry-DDX21-V5, and pUC19 HTLV-1 PBS mutant constructs

| Mutant | | Primer sequence |
|---|---|---|
| **pET3xc HTLV-1 Gag** | **ΔC29** | Forward primer: 5'-GAA AAC CTT TAC TTC CAG GGC GAT TAC G-3'<br>Reverse primer:5'- GAA GTA AAG GTT TTC AAT GGT TGG CTT CAG-3' |
| **pUCOgs HTLV-1 Gag-GFP** | **ΔMA** | Forward primer: 5'-CAA GCT TGC CAC CAT GCC TGT CAT GCA CC-3'<br>Reverse primer: 5'-CAT GGT GGC AAG CTT GAG CTC GAG ATC TG-3' |
| | **ΔCA** | Forward primer: 5'-GTG GTG CAG CCT AAA AAG CCC CCC CCG AAT C-3'<br>Reverse primer: 5'-TAG GCT GCA CCA CCA GCA CTT GGG GGG CGG-3' |
| | **ΔNC** | Forward primer: 5'-CAA GAC CAA GGT GCT GGG ATC CCT TGA AGT G-3'<br>Reverse primer: 5'-CAG CAC CTT GGT CTT GTC CTT GGG GGT CCA G-3' |
| | **ΔZF1** | Forward primer: 5'-CAC AGC CCA GAC CTC CCC CAG GAC CAT GC-3'<br>Reverse primer: 5'-GAG GTC TGG GCT GTG TGG GCT GAT TCG GGG-3' |
| | **ΔZF2** | Forward primer: 5'-CCA AGA CTG AAG CCA ACC ATT CCT GAG CC-3'<br>Reverse primer: 5'-GGC TTC AGT CTT GGT GGT CCT GGG GGA GGT C-3' |
| | **ΔNC CTD (ΔC29)** | Forward primer: 5'-CTG AAG CCA ACC ATT GGA TCC CTT GAA GTG-3'<br>Reverse primer: 5'-AAT GGT TGG CTT CAG TCT TGG GCA GTC GC-3' |
| **pUCOgs HTLV-1 Gag-GFP or Gag-FLAG** | **G2A** | Forward primer: 5'-ATG GCC CGC ATT TTC AGC CGC TCC GCT TC-3'<br>Reverse primer: 5'-GAA AAT GCG GGC CAT GGT GGC AAG CTT GAG-3' |
| | **M147A** | Forward primer: 5'-GGC AGG CCA AGG ATC TCC AGG CTA TTA AGC-3'<br>Reverse primer: 5'-GAT CCT TGG CCT GCC AAG GGC GGT GGT TG-3' |
| | **F177AQ178A** | Forward primer: 5'-CGT GCA AGC TGC TGA TCC AAC CGC TAA GG-3'<br>Reverse primer: 5'-CAG CAG CTT GCA CGG CCA GCC TAA TTG TC-3' |
| | **Y191A** | Forward primer: 5'-CAG GCC CTT TGC TCC AGC CTC GTC GCC AGC-3' |

| | | |
|---|---|---|
| | | Reverse primer: 5'-GGA GCA AAG GGC CTG CAG CAG GTC CTG GAG-3' |
| **pPB RPL7** | **ΔbZIP (ΔN77)** | Forward primer: 5'-GCT GGC AAC TTC TAT GTA CCT GCA GAA CC-3'<br>Reverse primer: 5'-CAT AGA AGT TGC CAG CGC CAA CTT TTT TG-3' |
| | **ΔCTD (ΔC51)** | Forward primer: 5'-GAG ATC TAT ACT GTT TGC CCA ACT TTC TTG-3'<br>Reverse primer: 5'-CAG TAT AGA TCT CAT GAA TCA AAT CCT CC-3' |
| **pCMV3 HA-RPL7** | **ΔbZIP (ΔN77)** | Forward primer: 5'-GCT GGC AAC TTC TAT GTA CCT GCA GAA CC-3'<br>Reverse primer: 5'-GAA GTT GCC AGC GCT ACC GCC TCC ACC-3' |
| | **bZIP (N77)-(GGGGS)$_3$-CTD (C51)** | Forward primer: 5'-GGT GGA GGA GGT AGC GGT GGT GGA GGA TCT GGA AAA CGC TTC AAA G-3'<br>Reverse primer: 5'-GCT ACC TCC TCC ACC AGA TCC ACC ACC TCC TTT TCT TGC CAT CCT C-3' |
| | **ΔCTD (ΔC51)** | Forward primer: 5'-GAG ATC TAT ACT GTT TGA ATC TAG AGC GGC-3'<br>Reverse primer: 5'-CAG TAT AGA TCT CAT GAA TCA AAT CCT CC-3' |
| | **bZIP (N77)** | Forward primer: 5'-GAG GAT GGC AAG AAA ATG AAT CTA GAG CGG C-3'<br>Reverse primer: 5'-CTT GCC ATC CTC GCC ATT CGA ATT TCA GTT C-3' |
| | **CTD (C51)** | Forward primer: 5'-GGA AAA CGC TTC AAA GAG GCA AAT AAC-3'<br>Reverse primer:5'-GAA GCG TTT TCC GCT ACC GCC TCC ACC-3' |
| **pLV CMV mCherry-DDX21-V5** | **ΔNTD** | Forward primer: 5'- GGC GCT TTC TCT AAT TTT CCC ATA TCT G-3'<br>Reverse primer: 5'- AGA GAA AGC GCC CAT GCT AGC TGA TCC G-3' |
| | **ΔHelicase N** | Forward primer: 5'- GTG GAA CAA AAA GAA GTG GAG CAT CTG GC-3'<br>Reverse primer: 5'- CTT TTT GTT CCA CAG GTA TTT C-3' |
| | **ΔHelicase C** | Forward primer: 5'- GAA AAC GGC AAT AAC TGC AAC AGA AAT AAT AAA AG-3'<br>Reverse primer: 5'- GTT ATT GCC GTT TTC TGA GTC-3' |
| | **ΔCTD** | Forward primer: 5'- GAA TAG GTG TTC CTT CTG CCT CGA GTC TAG AG-3'<br>Reverse primer: 5'- GAA GGA ACA CCT ATT CGT TTG-3' |
| | **ΔHelicase core** | Forward primer: 5'- GTG GAA CAA AAA GAA GCA ACA GAA ATA ATA AAA G-3'<br>Reverse primer: 5'- CTT TTT GTT CCA CAG GTA TTT C-3' |
| **pUC19 HTLV-1 PBS** | **Δ425-434** | Tailed forward primer: 5'-GGC TCG TCC GGG ATA TTT ATT CCC TAG GCA ATG GG-3'<br>Forward primer: 5'-TTT ATT CCC TAG GCA ATG GG-3'<br>Tailed reverse primer: 5'-TAT CCC GGA CGA GCC CCC AAC TGT GTA CTA AAT TTC TC-3'<br>Reverse primer: 5'-CCC AAC TGT GTA CTA AAT TTC TC-3' |

**Table S2.** List of RNA sequences used in this study

| RNA | Length (nt) | Sequence |
|---|---|---|
| HTLV-1 WT PBS[α] | 98 | 5'-*GG*G CCC AUC CUA UAG CAC UCU CCA GGA GAG AAA UUU AGU ACA CAG UUG GGG GCU CGU CCG GGA UAC GAG CGC CCC UUU AUU CCC UAG GCA AUG GGC C*C*-3' |
| HTLV-1 Δ425-434 PBS[α] | 88 | 5'-*GG*G CCC AUC CUA UAG CAC UCU CCA GGA GAG AAA UUU AGU ACA CAG UUG GGG GCU CGU CCG GGA UAU UUA UUC CCU AGG CAA UGG GCC *C*-3' |
| $tRNA^{Pro}_{UGG}$ | 75 | 5'-GGC UCG UUG GUC UAG GGG UAU GAU UCU CGC UUU GGG UGC GAG AGG UCC CGG GUU CAA AUC CCG GAC GAG CCC CCA-3' |

[α] The sequences underlined are the HTLV-1 18-nt PBS. Italicized 5' 2 G's are not encoded by HTLV-1 and were added for efficient initiation of *in vitro* transcription by T7 RNA polymerase.

**Table S3.** List of all AP-MS protein hits

| Gene symbol | Protein name | Total Spectrum Count | |
|---|---|---|---|
| | | Gag-GFP | GFP |
| RPN1 | Dolichyl-diphosphooligosaccharide--protein glycosyltransferase subunit 1 | 50 | 4 |
| RPL4 | 60S ribosomal protein L4 | 48 | 1 |
| ABCD3 | ATP-binding cassette sub-family D member 3 | 37 | 4 |
| RPL7A | 60S ribosomal protein L7a | 34 | 2 |
| RPL6 | 60S ribosomal protein L6 | 31 | 0 |
| **RPL7** | **60S ribosomal protein L7** | 25 | 0 |
| RPS3A | 40S ribosomal protein S3a | 23 | 3 |
| **DDX21** | **Nucleolar RNA helicase 2** | 22 | 0 |
| RPL18 | 60S ribosomal protein L18 | 20 | 0 |
| DDOST | Dolichyl-diphosphooligosaccharide--protein glycosyltransferase 48 kDa subunit | 20 | 1 |
| RPLP2 | 60S acidic ribosomal protein P2 | 20 | 3 |
| RPL12 | 60S ribosomal protein L12 | 19 | 0 |
| RPS8 | 40S ribosomal protein S8 | 19 | 0 |
| ATG2A | Autophagy-related protein 2 homolog A | 19 | 0 |
| RPN2 | Dolichyl-diphosphooligosaccharide--protein glycosyltransferase subunit 2 | 18 | 0 |
| MCU | Calcium uniporter protein, mitochondrial | 18 | 1 |
| LBR | Lamin-B receptor | 18 | 3 |
| NMT1 | Glycylpeptide N-tetradecanoyltransferase 1 | 17 | 0 |
| RPL10A | 60S ribosomal protein L10a | 17 | 1 |
| RPL15 | 60S ribosomal protein L15 | 16 | 0 |
| GNL3 | Guanine nucleotide-binding protein-like 3 | 16 | 1 |
| RPS6 | 40S ribosomal protein S6 | 16 | 2 |
| MYBBP1A | Myb-binding protein 1A | 15 | 0 |
| RPL23A | 60S ribosomal protein L23a | 14 | 0 |
| RPL8 | 60S ribosomal protein L8 | 14 | 0 |
| DHX9 | ATP-dependent RNA helicase A | 14 | 3 |
| RPL27 | 60S ribosomal protein L27 | 13 | 0 |
| TMEM43 | Transmembrane protein 43 | 13 | 1 |
| LMAN2 | Vesicular integral-membrane protein VIP36 | 12 | 0 |
| ALG8 | Probable dolichyl pyrophosphate Glc1Man9GlcNAc2 alpha-1,3-glucosyltransferase | 12 | 0 |
| DGAT1 | Diacylglycerol O-acyltransferase 1 | 11 | 0 |
| NOP2 | Probable 28S rRNA (cytosine(4447)-C(5))-methyltransferase | 11 | 0 |
| NOM1 | Nucleolar MIF4G domain-containing protein 1 | 11 | 0 |
| GPC4 | Glypican-4 | 10 | 0 |
| HSPA4 | Heat shock 70 kDa protein 4 | 10 | 0 |

| | | | |
|---|---|---|---|
| BCAP31 | B-cell receptor-associated protein 31 | 10 | 0 |
| ABCB10 | ATP-binding cassette sub-family B member 10, mitochondrial | 10 | 0 |
| NOP56 | Nucleolar protein 56 | 10 | 1 |
| RBM4 | RNA-binding protein 4 | 9 | 0 |
| SCAMP3 | Secretory carrier-associated membrane protein 3 | 9 | 0 |
| PEX10 | Peroxisome biogenesis factor 10 | 9 | 0 |
| SON | SON DNA binding protein | 9 | 0 |
| PTDSS1 | Phosphatidylserine synthase 1 | 9 | 0 |
| PIGK | GPI-anchor transamidase | 9 | 0 |
| RRBP1 | Ribosome-binding protein 1 | 9 | 0 |
| SEC61A1 | Protein transport protein Sec61 subunit alpha isoform 1 | 9 | 1 |
| SEC61A2 | Protein transport protein Sec61 subunit alpha isoform 2 | 8 | 0 |
| RBM4B | RNA-binding protein 4B | 8 | 0 |
| RPLP0 | 60S acidic ribosomal protein P0 | 8 | 0 |
| MLEC | Malectin | 8 | 0 |
| STEAP3 | Metalloreductase STEAP3 | 8 | 0 |
| DHX30 | Putative ATP-dependent RNA helicase DHX30 | 8 | 0 |
| RPL34 | 60S ribosomal protein L34 | 8 | 1 |
| SAAL1 | Synoviocyte proliferation-associated in collagen-induced arthritis protein 1 | 8 | 1 |
| TUBGCP2 | Gamma-tubulin complex component 2 | 8 | 1 |
| YBX1 | Nuclease-sensitive element-binding protein 1 | 7 | 0 |
| CAND2 | Cullin-associated NEDD8-dissociated protein 2 | 7 | 0 |
| CYB5R3 | NADH-cytochrome b5 reductase 3 | 7 | 0 |
| RPL35A | 60S ribosomal protein L35a | 7 | 0 |
| SOAT1 | Sterol O-acyltransferase 1 | 7 | 0 |
| RPL18A | 60S ribosomal protein L18a | 7 | 0 |
| EMC1 | ER membrane protein complex subunit 1 | 7 | 0 |
